# The prevalence of motility within the human oral microbiota

**DOI:** 10.1101/2023.07.17.549387

**Authors:** Sofia T. Rocha, Dhara D. Shah, Qiyun Zhu, Abhishek Shrivastava

## Abstract

The human oral and nasal microbiota contains approximately 770 cultivable bacterial species. More than 2000 genome sequences of these bacteria can be found in the expanded Human Oral Microbiome Database (eHOMD). We developed HOMDscrape, a freely available Python software tool to programmatically retrieve and process amino acid sequences and sequence identifiers from BLAST results acquired from the eHOMD website. Using the data obtained through HOMDscrape, the phylogeny of proteins involved in bacterial flagellar motility, Type 4 pilus driven twitching motility, and Type 9 Secretion system (T9SS) driven gliding motility was constructed. A comprehensive phylogenetic analysis was conducted for all components of the rotary T9SS, a machinery responsible for secreting various enzymes, virulence factors, and enabling bacterial gliding motility. Results revealed that the T9SS outer membrane ß-barrel protein SprA of human oral microbes underwent horizontal evolution. Overall, we catalog motile microbes that inhabit the human oral microbiota and document their evolutionary connections. These results will serve as a guide for further studies exploring the impact of motility on shaping of the human oral microbiota.

## BACKGROUND

The human oral cavity has different parts that are colonized by microbes. These include, but are not limited to, teeth, tongue, cheek, nose, and gingival plaques (1). The human oral microbiota is linked to diseases such as caries (2, 3), tonsillitis (4), gingivitis, periodontitis (5, 6), cardiovascular diseases (7, 8), oral cancer (9, 10), and Alzheimer’s disease (11, 12). Caries occurs when acid-producing bacteria break down dental enamel and dentine, leading to mineral dissolution (3, 13). Bacterial tonsillitis is characterized by inflammation of the tonsils (4, 14) while gingivitis causes red, swollen, and bleeding gums (13). Periodontitis is a more severe form of inflammation that results in gum and bone deterioration and subsequent bone loss. Both gingivitis and periodontitis are associated with the formation of polymicrobial biofilms (5, 6). The oral pathogen *Porphyromonas gingivalis* uses the Type 9 Secretion system (T9SS) to secrete harmful gingipain proteases. These proteases affect the immune system and cause periodontitis (15, 16). Gingipains also have neurotoxic properties and are associated with Alzheimer’s disease (11, 12) and cardiovascular diseases (17–20).

The oral cavity has different gradients influenced by factors like nutrient availability, heat, oxygen, saliva, and crevicular fluid (21). Bacteria that might possess the ability to sense some of these gradients could move towards favorable conditions using chemotaxis. This directed movement may influence the micron-scale spatial distribution of the human oral microbiota. Bacteria employ various motility mechanisms, with flagellar, twitching, and gliding motility being the most notable modes of motility (22–24). Many pathogens and commensals display robust motility. Here, we catalog motile microbes found in the human oral cavity that are driven by flagellar, twitching, and gliding motility.

Flagellated bacteria have a rotating flagellar bundle (FliC) driven by a motor powered by a proton motive force. Regulated by the chemotaxis pathway, the flagellar motor enables the bacteria to swim in a specific direction (run). Disruption in the direction of the flagellar motor results in a tumble. The probability of running or tumbling depends on the chemotaxis gradient (24). Twitching motility is a type of surface-associated movement facilitated by Type 4 pili. Slender filaments (PilA, PilE) adhere to a solid surface and undergo rapid extension cycles at their ends. The filament retraction, powered by an ATP-driven motor called PilT, pulls the cell forward (25–27). In contrast, the bacterial T9SS, which is a recently discovered machinery, enables both gliding motility and protein secretion. It utilizes a molecular rack and pinion machinery for gliding, where the T9SS rotary motor acts as a pinion to move a conveyor belt rack (GldJ) on the bacterial cell surface. This system facilitates the movement of adhesins along the cell surface (23, 28). Motile T9SS containing human oral microbes carry other oral microbes and bacteriophages as cargo (29). A functional T9SS allows for the secretion of over 200,000 bacterial proteins in members of the Bacteroidetes-Fibrobacteres-Chlorobi superphylum (23).

The Human Oral Microbiome Database (eHOMD) houses over 2000 genome sequences from human oral and nasal isolates, encompassing 775 bacterial types across 185 genera (30). In this study, an extensive search was conducted on eHOMD to identify proteins involved in the three motility systems described above. Computational tools were developed to automate the retrieval of sequence identifiers and amino acid sequences for the identified proteins. This resulted in a comprehensive catalog of motile microbes found in the human oral microbiota. A phylogenetic analysis shed light on the evolutionary lineages of these motility-related proteins. Interestingly, the analysis of the T9SS machinery, which is relatively understudied, revealed evidence of horizontal gene transfer in the outer-membrane pore protein and regions of structural similarities between T9SS rotor and conveyor belt proteins. The results presented here not only enabled the cataloging of motile bacteria present in the oral microbiota but also provides insight into their evolutionary relationships.

## RESULTS

### Development and use of HOMDscrape for automated generation of a curated dataset from BLAST results on eHOMD

The version 2 of HOMD releases all their BLAST results with a sequence identifier and does not have a user-friendly function that allows a user to save sequence identifiers along with the species name and amino acid sequence. To automatically create a database that can be used for further analysis, a local software named HOMDscrape was developed to programmatically retrieve and process eHOMD search results.

HOMDscape is a Python-based software tool freely available on GitHub under an MIT license (https://github.com/strocha1/HOMDscrape). It utilizes the chromedriver library (version 96.0.4664.45) to automate interaction with the eHOMD webserver using an open-source API and the HTML code from eHOMD. It is used to automate the process of gathering species names and amino acid sequences from the BLAST result. A schematic depicting each step of HOMDscrape is outlined in **Figure 1**. Once the raw data is collected by HOMDscrape, it is converted into a FASTA format and saved as a plain text file that can be used for downstream analysis.

**Figure 1.**
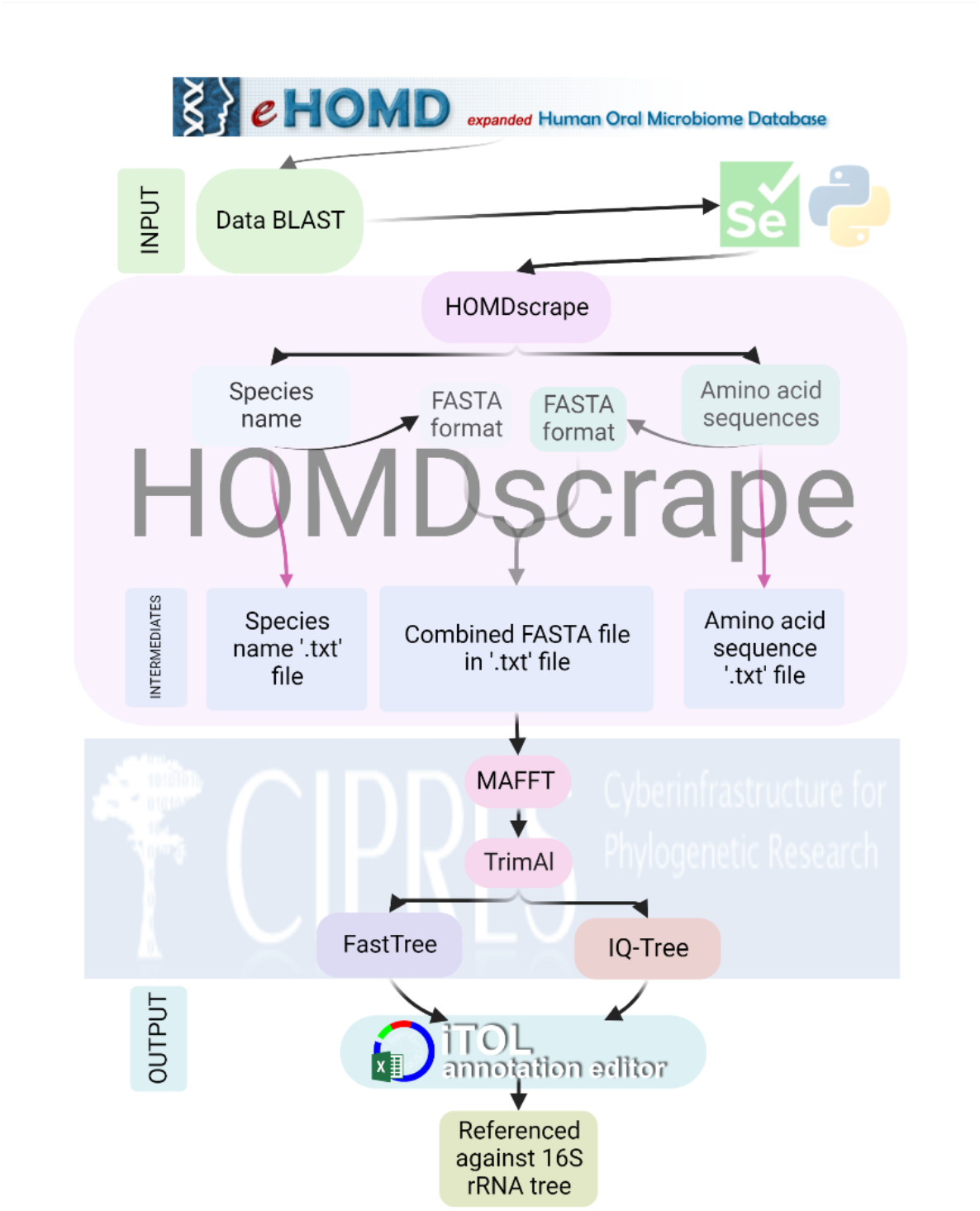
An outline of HOMDscrape, a selenium Python software that extracts bacterial species names and amino acid sequences from BLAST results on eHOMD.

### A phylogenetic tree of human oral microbes cataloged on eHOMD

The phylogenetic tree of eHOMD-hosted genomes was reconstructed using their 16S rRNA gene sequences **(Figure S1).** This tree illustrates the evolution of all the species described on the eHOMD. The 16S rRNA RefSeq data available on eHOMD was used as raw data for phylogenetic reconstruction. The trees constructed later in this study, with the data collected by HOMDscrape, were compared with this 16S tree. Except for this 16S rRNA analysis, all phylogenetic analyses were conducted using the data obtained and parsed by HOMDscrape.

### Prevalence of flagellar motility in the human oral microbiota

The flagellar motor and the virulence-associated type III injectosome share several structural proteins (31, 32) and both systems have the type III secretion system for protein export. To reduce false positives during our bioinformatics search for flagellar motility, it was important to exclude flagellar proteins that are found in the type III secretion system or the type III injectosome. We compared the proteins that form the flagellar motor and the type III injectosome **(Table S1)**. This comparison led to the selection of three proteins, FliC, FlgK, and FlgL as markers for the presence of flagellar motility. FliC is the main structural subunit in the flagellar filament. FlgK and FlgL form the hook-filament junction (31, 33).

We predict that 47 genera in the human oral microbiota have flagellar motility **(Figure 2).** We identified FliC in 84 species of 48 genera, FlgK in 108 species of 64 genera, and FlgL in 75 species of 49 genera **(Figure 2 and Table S2).** The evolutionary pattern of FliC, FlgK, and FlgL in human oral microbes follows the 16S rRNA tree **(Figure 2 and Figure S1)**. For all IQ Trees (34) presented in this study, the periphery of the tree is labelled by a colored circle that represents the genus corresponding to the terminal node. The branches are colored corresponding to the bootstrap value. The colormap indicates bootstrap values.

**Figure 2.**
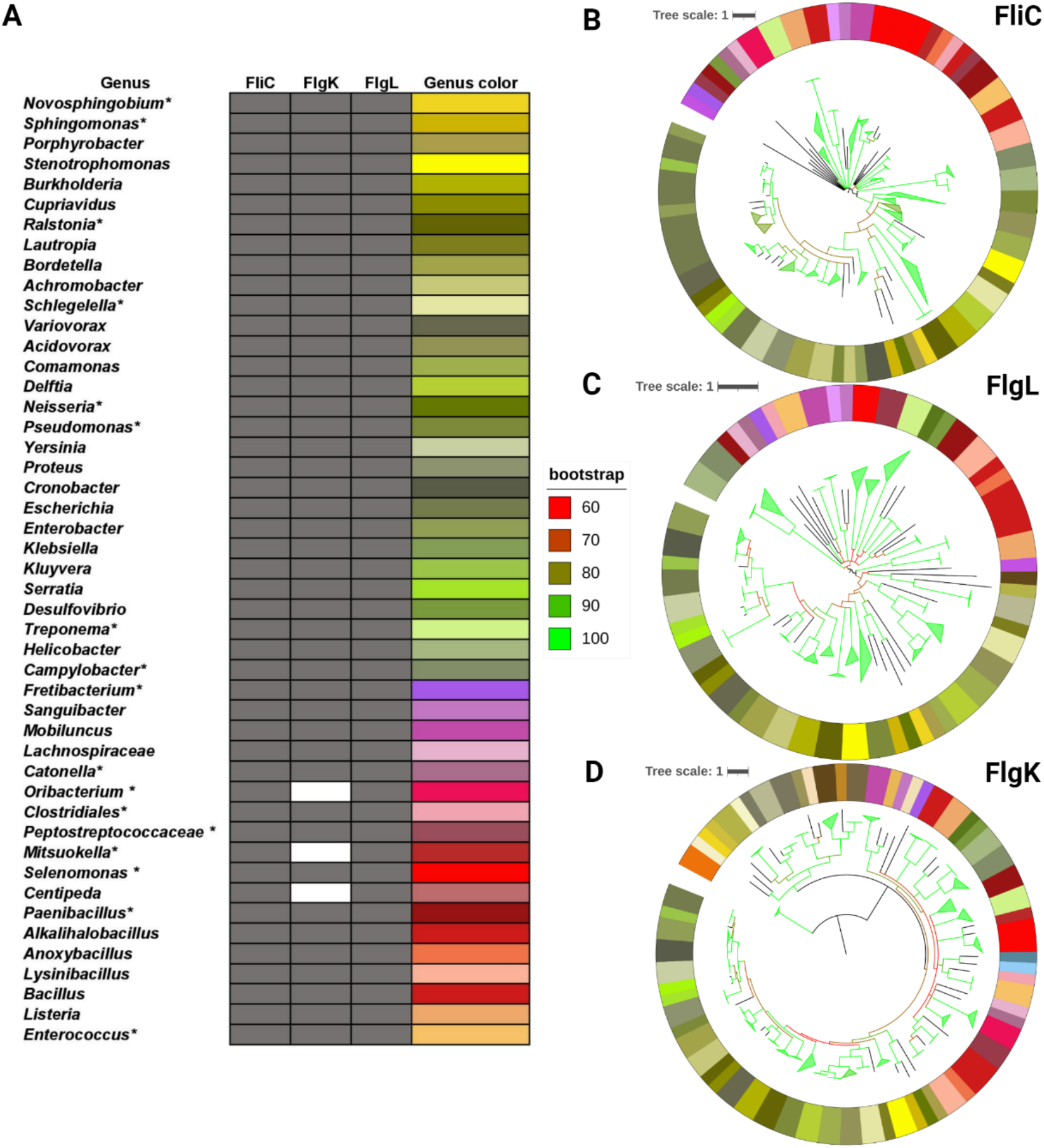
**A.** Summary of the genera containing flagellar motility proteins. Genera marked with ‘*’ display differences at the species level. A comprehensive list with species names is available in Table S2. **B-D.** IQ Trees of FliC, FlgL and FlgK. The colormap for IQ Trees is depicted via the column ‘Genus color’ in Fig. 2A.

### Prevalence of type 4 pilus driven twitching motility in the human oral microbiota

The type 4 pilus motor shares several homologous proteins with the type 2 secretion system (T2SS) (25). Proteins that were found in both type 4 pilus machinery and the T2SS were excluded from the downstream analysis and three structural proteins, PilT, PilE, and PilA, were used as markers for the presence of Type 4 pilus driven twitching motility. PilT functions as the retraction ATPase that aids in twitching motility. PilE functions as the major pilin in bacteria such as *Neisseria gonorrhoeae* and a minor pilin in *Pseudomonas aeruginosa* while PilA functions as a major pilin in *P. aeruginosa* (25). We predict that 32 genera in the human oral microbiota have twitching motility **(Figure 3).** PilT is found in 105 species of 56 genera, making up to 30% of the total genera found in the human oral microbiota. In addition, 76 species of 30 genera had PilE and 85 species of 35 genera had PilA **(Figure 3 and Table S3)**. The evolutionary pattern of PilT, PilE, and PilA in human oral microbes follows the 16S rRNA tree **(Figure 3 and Figure S1)**.

**Figure 3.**
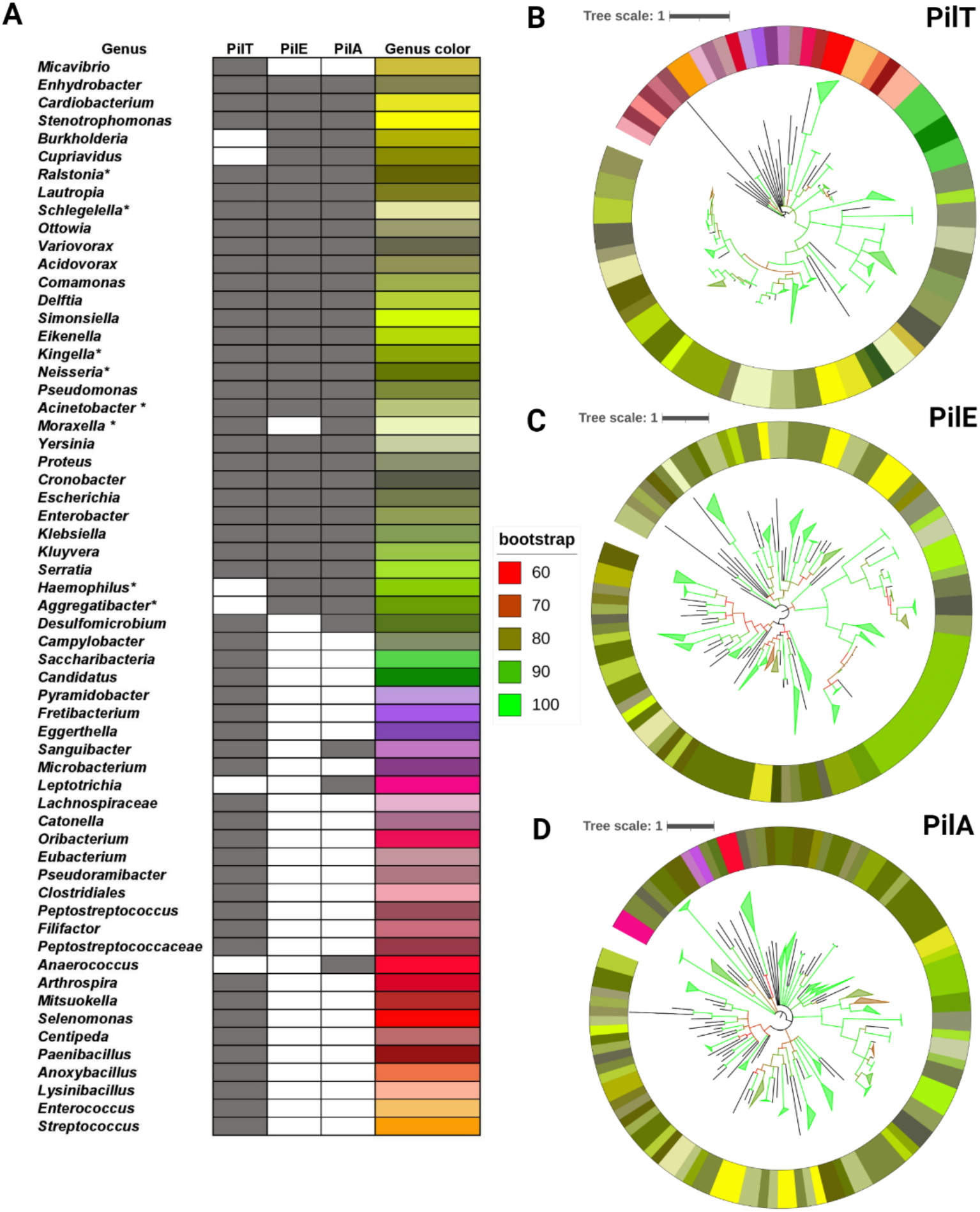
**A.** Summary of the genera containing Type 4 pili twitching motility proteins. Genera marked with ‘*’ display differences at the species level. A comprehensive list with species names is available in Table S2. **B-D.** IQ Trees of PilE, PilT, and PilA. The colormap for IQ Trees is depicted via the column ‘Genus color’ in Fig. 3A.

### Prevalence of T9SS in the human oral microbiota

T9SS has several structural proteins that are necessary for both secretion and motility, while some structural proteins are only involved in motility (23). T9SS is a recently discovered machinery and very little is known about its evolution. To fill this knowledge gap, we searched for 16 T9SS proteins and analyzed their evolutionary patterns. We predict that a functional T9SS is found in 68 species from 7 genera are isolated from the human oral microbiota **(Figure 4).**

**Figure 4.**
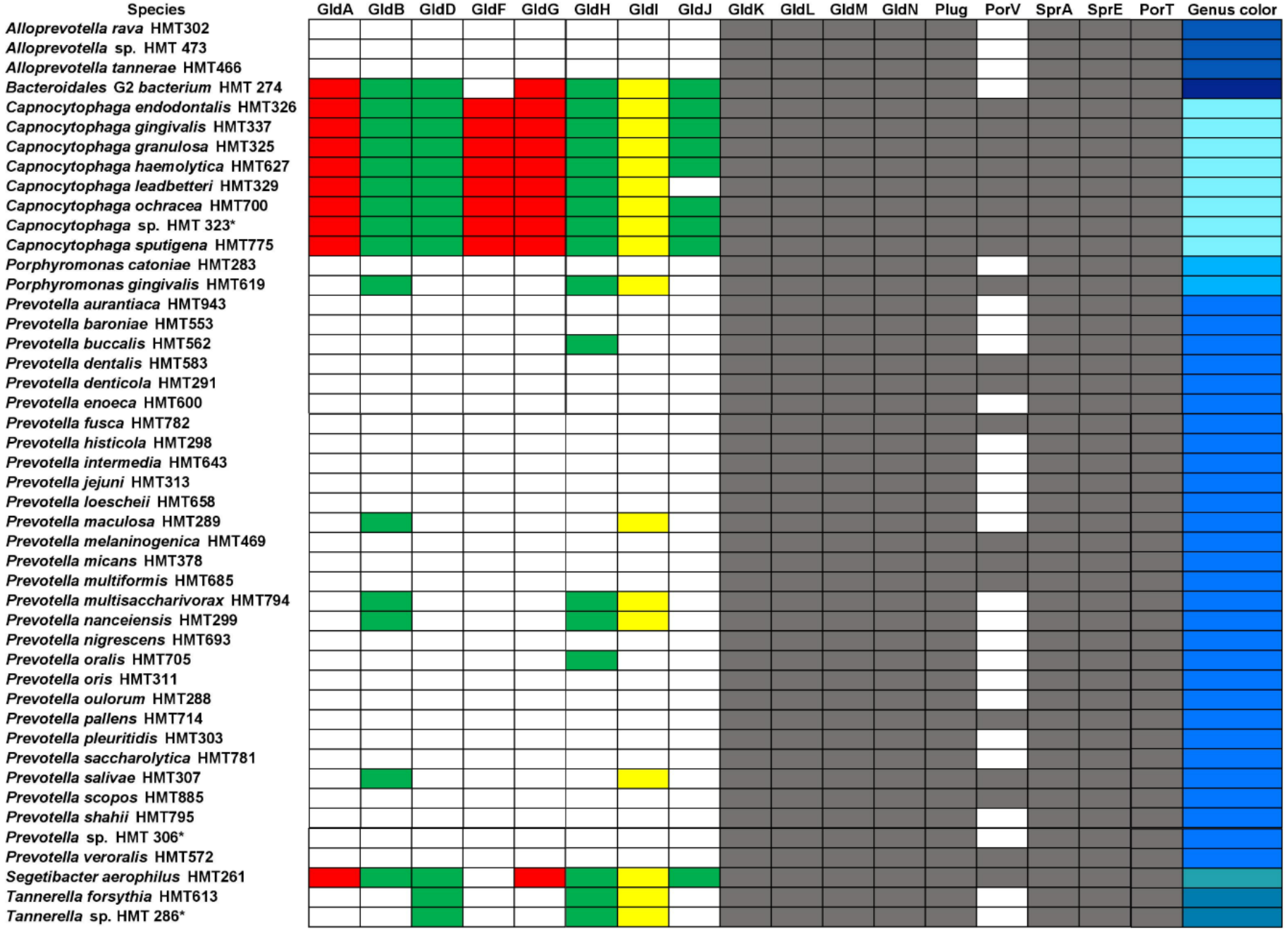
Summary of the genera containing Type 9 gliding motility proteins. Species marked with a ‘*’ represent unnamed isolates. A comprehensive list with species names is available in Table S2.

The predicted rotary component of T9SS is made up of four proteins. GldK/PorK, GldL/PorL, GldM/PorM, and GldN/PorN. GldK and GldN form a periplasmic ring, while GldL and GldM form a transmembrane ion channel that powers T9SS rotation (23, 35, 36). GldL and GldM are structurally similar to the 5:2 MotA-MotB complex found in the flagellar motor. However, there is no sequence similarity between GldL-GldM and MotA-MotB (36). Using HOMDscrape, we found that 71 species of oral microbes belonging to seven genera contained GldK. Additionally, 70 species within the same seven genera had GldL, while 71 species had GldM, and 68 species had GldN **(Figure 5)**. In summary, our analysis showed that seven different genera of eHOMD cataloged microbes, namely *Capnocytophaga, Porphyromonas*, *Prevotella, Bacteroidales [G-2], Tannerella, Segetibacter* and *Alloprevotella* possess the core components of T9SS **(Figure 4)**.

**Figure 5.**
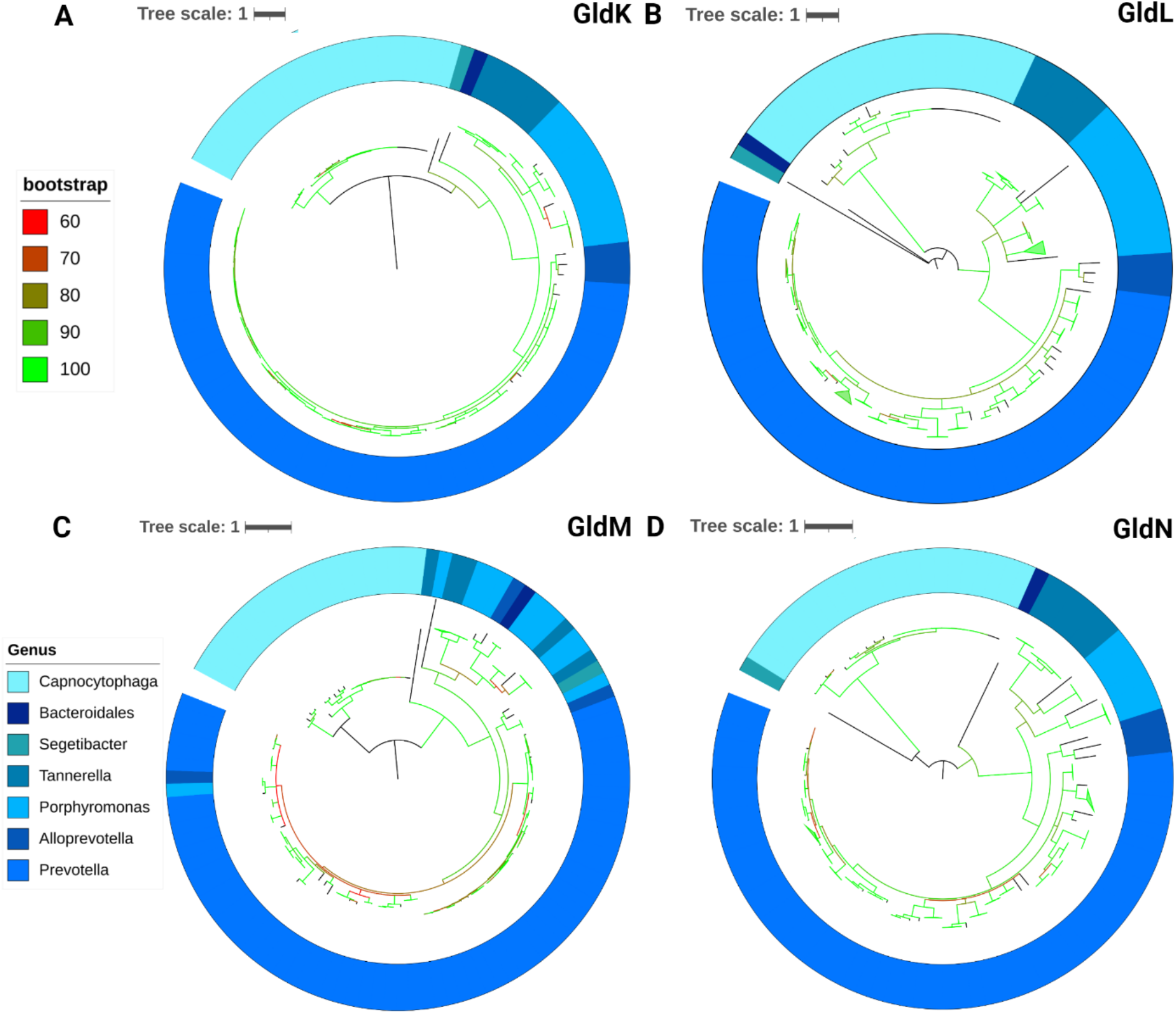
**A-D.** IQ Trees of GldK, GldL, GldM, and GldN where the colormap is depicted via the column ‘Genus color’ in Fig. 4.

PorT (SprT) is an outer membrane protein that is required for the secretion of gingipains in *P. gingivalis* (23, 37, 38). We found that 71 oral microbial species within nine genera had PorT **(Figure S2)**. SprE (PorW) is required for the secretion for a variety of T9SS proteins and is predicted to act as a bridge between the SprA translocon and GldK/N rings (23, 35). We found that 92 oral microbial species within 19 genera had SprE **(Figure 4, Figure S2)**.

### Horizontal evolution of the T9SS translocon

SprA is the largest single polypeptide outer membrane ß-barrel that acts as the protein translocon of T9SS (23). SprA interacts with PorV and the T9SS Plug to form an outer membrane gate (39). We found that 73 oral microbial species within nine genera had SprA **(Figure 6).** 41 oral microbial species within nine genera had PorV, and 72 oral microbial species within nine genera had the T9SS Plug. Members of the *Melioribacter* and *Ignavibacterium* genera contain the PorV, SprA, and Plug complex but does not contain any other T9SS protein **(Figure 6, Figure S3)**.

**Figure 6.**
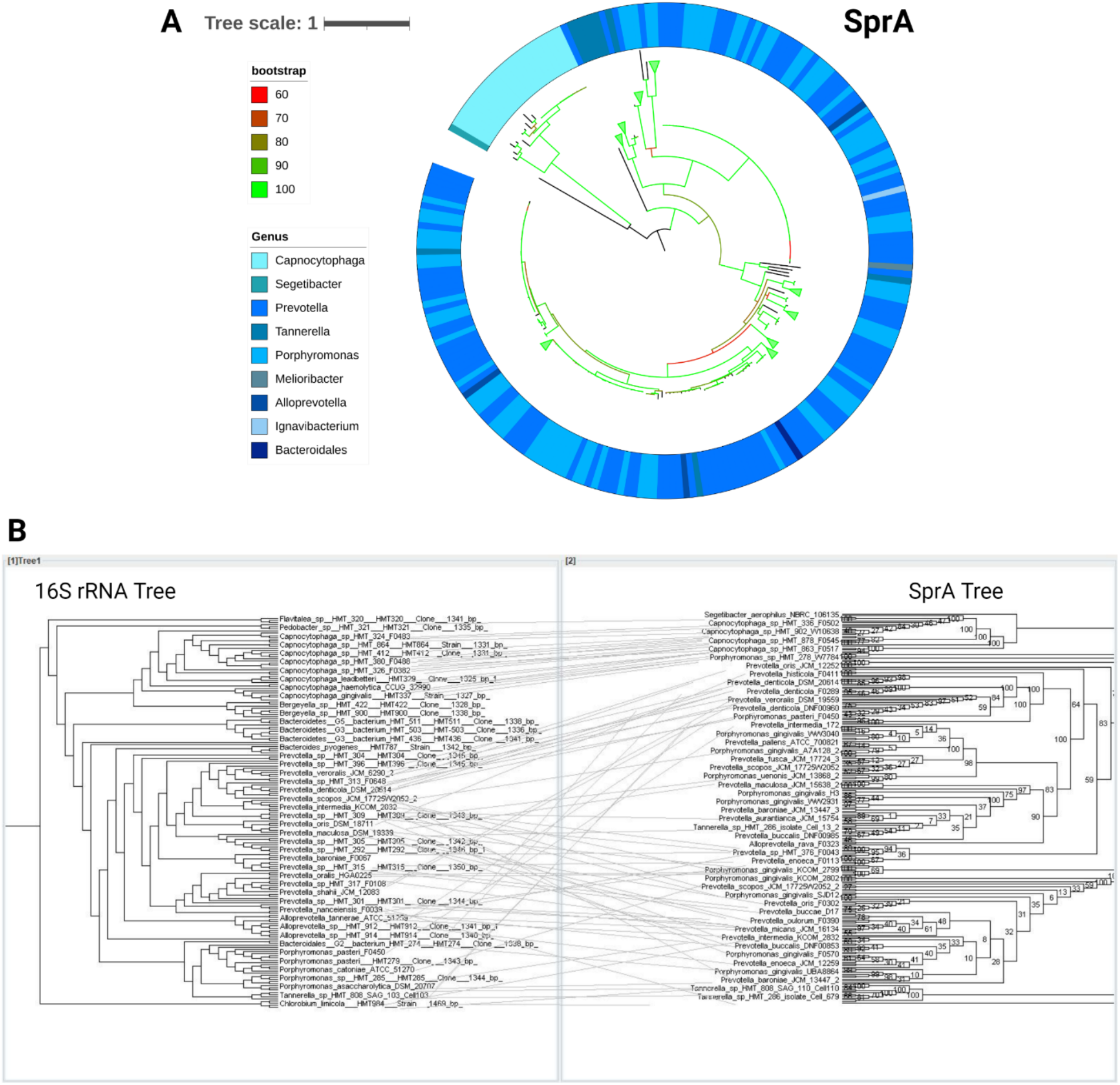
**A.** IQ Tree of SprA where the colormap is depicted via the column ‘Genus color’ in Fig. 4. **B.** Tanglegram of 16S rRNA tree (Left) and SprA (Right).

The phylogenetic tree of SprA had a high volume of inconsistent terminal species when compared to the phylogenetic tree of the 16S rRNA gene **(Figure 6)**. The terminal nodes were inconsistent after the *Capnocytophaga* genus. This suggests that there was a higher level of recombination between other genera containing SprA. Thus, there is a possibility that the SprA protein is not well conserved, causing the deviation from the 16S rRNA phylogenetic tree. SprA is a 267 KDa polypeptide and has roughly 7 domains (39). In order to determine if the deviation from the 16S rRNA tree is found throughout the whole protein or is limited to certain domains, a series of tanglegrams were constructed for each SprA domain **(Figure S4-S10)** (40). While there was deviation from the 16S rRNA tree across all domains, a lower rate of deviation was found in domains 2, 5, and 7 as compared with the other four domains **(Figure S5, S8, S10)**.

### Prevalence and phylogeny of the ABC transporter domain containing gliding proteins

GldA is essential for gliding motility, and it is predicted to be an ABC transporter that interacts with T9SS. GldG is predicted to be a transmembrane protein that interacts with GldF and GldA to form an ABC transporter. GldF is essential for gliding motility and is required for the stability of GldK. Whereas GldB and GldD are lipoproteins that are essential for gliding motility (41, 42). Using HOMDscrape, 22 oral microbial species in three genera were discovered to contain GldA. In parallel, 29 oral microbial species in seven genera had GldB and 25 oral microbial species in four genera contained GldD and 22 oral microbial species in three genera had GldF **(Figure S11)**. Similarly, 22 oral microbial species in three genera had GldG **(Figure 12)**.

### Phylogeny of proteins that either form or stabilize the mobile cell-surface track driven by T9SS

GldH and GldI are essential for gliding motility and help stabilize GldJ (23, 43, 44). 39 oral microbial species in eight genera had GldH. 32 oral microbial species in eight genera had GldI. However, only 21 oral microbial species in three genera had GldJ **(Figure 12)**.

GldJ forms helical tracks on the cell surface that acts as the rack in the rack and pinion assembly that drives gliding motility (41, 45). When comparing the phylogenetic tree of the T9SS gliding protein GldJ to other core T9SS proteins, a loss of genera is observed **(Figure 7D).** GldJ containing motile *Capnocytophaga* sp. constitute around 1% of the total diversity of the human oral microbiota but constitute 10% of total microbial abundance at important gingival sites within the human oral cavity (30, 46, 47).

**Figure 7.**
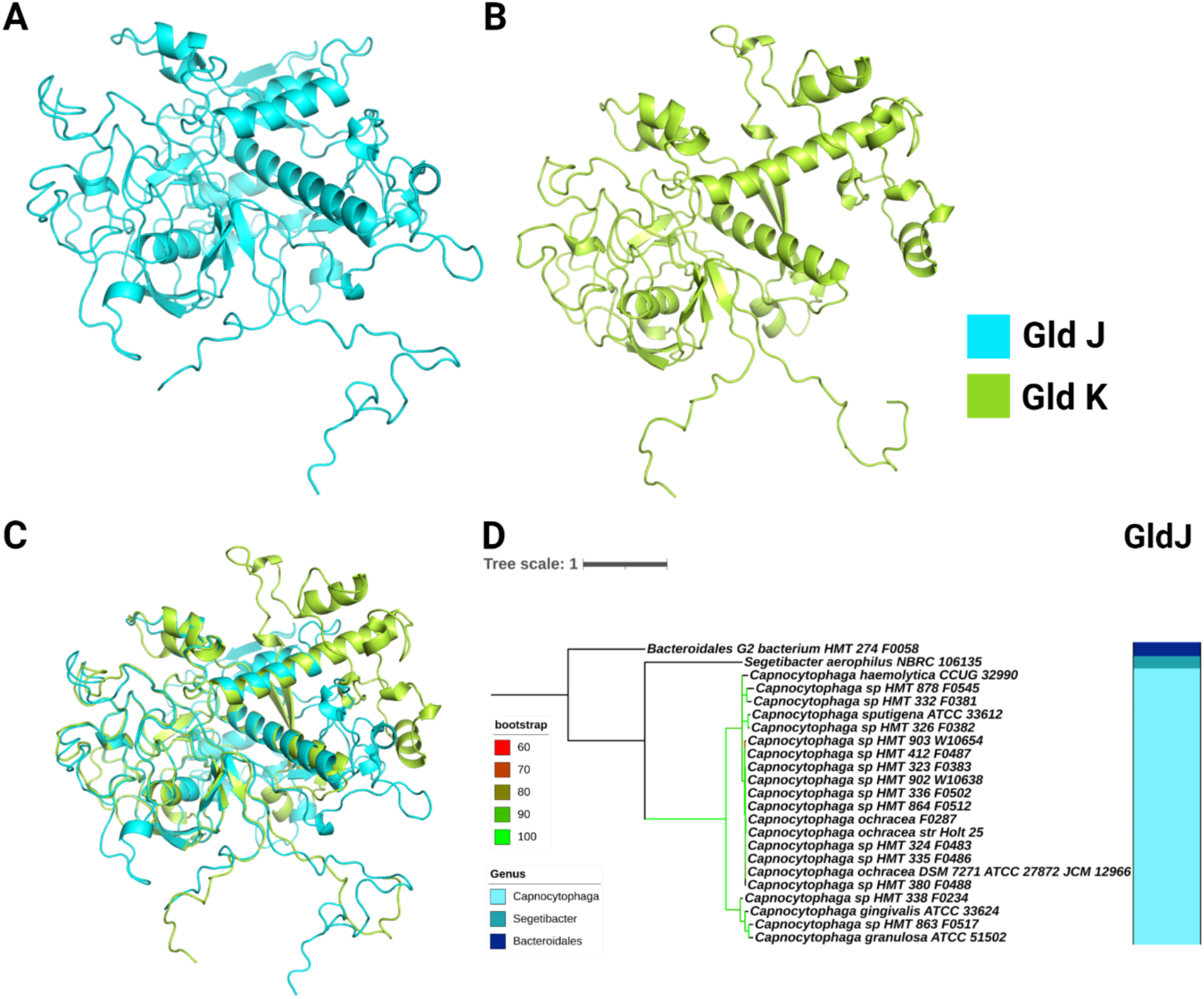
**A.** The predicted structure of GldJ. **B.** The predicted structure of GldK. **C.**

### Similarity between GldJ and GldK

GldJ of *Capnocytophaga gingivalis* has 31% amino acid sequence similarity to GldK of *Capnocytophaga gingivalis*. GldJ and GldK are lipoproteins that exhibit sequence similarity to each other and also to sulfatase-modifying enzymes such as formylglycine-generating enzyme (FGE), but lack certain active site residues of FGE (48).

The predicted Alphafold (49) structures of GldJ and GldK were generated **(Figure 7)** and the confidence in predictions per residue through the local distance difference test score (lDDT) (50) was plotted against residue positions (**Figure S13)**. Aligned GldJ and GldK show RMSD of 0.58 Å for the total of 1371 atoms. At the N-terminus of both proteins, beta sheets consisting of residues 39-42 of GldJ and 25-28 of GldK, 45-49 of GldJ and 31-35 of GldK, 65-69 of GldJ and 51-55 of GldK superimposed completely. Additionally alpha helices at the N-terminus with residues spanning 81-94 of GldJ and 67-82 of GldK were also well aligned in both structures. In the middle part of the protein, residues 148-158 of GldJ and 251-261 of GldK, and residues 238-246 of GldJ and 285-293 of GldK that form alpha helices, show complete alignment. At the C-terminus of both proteins, there are 3 beta sheets that are completely aligned. These are residues 487-490 of GldJ and 384-387 of GldK, residues 508-511 of GldJ and 405-408 of GldK, and residues 523-526 of GldJ and 420-423 of GldK. All residues are numbered after removal of the signal peptides at the N-terminus of both GldJ and GldK.

## DISCUSSION

Phylogenetics extensively utilizes the 16S rRNA gene to compare and classify organisms based on their evolutionary relationships. This method assumes that the gene is only vertically inherited and is not exchanged between different clades (51–53). Thus, it is used as a marker gene for evolutionary comparison and classification of bacteria. The terminal species shown on the phylogenetic trees of the motility proteins are expected to match the terminal species on the phylogenetic tree of the 16S rRNA gene. In most cases, this was true for the phylogenetic trees of each motility protein component, with the exception of the T9SS protein SprA. The IQ Trees for the T9SS protein component GldM, Type 4 Twitching motility protein PilA, and flagellar motility FlgK exhibited minor inconsistencies that may be due to annotation errors. In the IQ Tree of the T9SS protein component SprA, there was a relatively higher volume of inconsistent terminal species when compared to the IQ Tree of the 16S rRNA gene, suggesting that the T9SS SprA may be able to jump across lineages **(Figure 6)**.

Organisms of the Bacteroidetes phylum play important roles in the human gut microbiota and several species can have a mutualistic behavior with some members of the phylum displaying an opportunistic pathogen behavior. Changes in their abundance correlates with diseases such as diabetes, obesity, and ulcerative colitis (54–57). Bacteroidetes in the human oral microbiota employ their T9SS for diverse purposes. For instance, *P. gingivalis* secretes immunomodulatory gingipain proteases through the T9SS, while *Tannerella forsythia* uses the T9SS to transport virulence-associated cargo proteins to their cell surfaces (58). In motile organisms such as *Capnocytophaga ochracea*, the T9SS enables gliding motility and biofilm formation (59). T9SS driven gliding bacteria such as *Capnocytophaga gingivalis* transport other non-motile oral microbial species as cargo and they play a potential role in shaping the biogeography of the human oral microbiota (29).

## CONCLUSIONS

Our analysis suggests that around 25% of the genera found in the human oral microbiota have flagellar motility. Additionally, 30% of the genera found in the human oral microbiota encoded the proteins necessary for type 4 pilus driven motility. The analysis presented here shows that although around 4% of the genera diversity found in the human oral microbiota encode the T9SS **(Figure 5)**, they constitute about 15% of total microbial abundance at important gingival sites within the human oral cavity (48, 49). The T9SS enhances adhesion to surfaces and other microorganisms, which could potentially aid in the improvement of colonization. Additionally, certain enzymes secreted by T9SS have the capability to break down complex polysaccharides and human tissue, making them a potential source of nutrients for a diverse microbial community. These characteristics may contribute to the proliferation of T9SS-containing microorganisms within the oral microbiota. The ability to move is conserved in several oral commensals and pathogenic bacterial species **(Figure 2, Figure 3, Table S2)** but very little is known about their role in shaping the human oral microbiota. Our analysis sheds light on the evolution of several proteins that enable motility and the secretion of virulence factors in the oral microbiota.

Overall, the catalog of motile human oral microbes described in this study could serve as a database for future research on the influence of motility in shaping the human microbiota. The HOMDscrape tool is accessible for free and was created to build a database of motile microorganisms found in the human oral microbiota. HOMDscrape is a generalized tool that is not only useful for studying motility but can easily be used for the analysis of other proteins and generalized bioinformatics tasks on eHOMD.

## METHODS

### Choice and search for genes encoding motility proteins in eHOMD

The following template amino acid sequences were searched using Basic Local Alignment Search Tool (BLAST) against the genomes hosted at eHOMD. The template sequences were derived from the following model organisms: Flagellar system – *Escherichia coli,* Type 4 pili system – *Neisseria meningitidis and Pseudomonas aeruginosa,* T9SS – *Capnocytophaga gingivalis*. BLAST hits were filtered depending on the protein’s known functions, query sequence similarity, and E-values (35). The E-values cutoffs used for each protein are described in **Figure S15**. New genomes and updated bacterial names can be added with newer versions of HOMD. The aim of this study is to be exhaustive in cataloging of motility proteins in the current version of HOMD. As HOMD gets updated, we call on the community to help update the future iterations of this catalog.

### Multiple sequence alignment

n order to create phylogenetic trees of each motility protein, the following programs were used from the Cyberinfrastructure for Phylogenetic Research computing cluster (CIPRES) (60): MAFFT (Multiple Alignment using Fast Fourier Transform) on Extreme Science and Engineering Discovery Environment (XSEDE) (7.490) (61), TrimAl on XSEDE (1.2.59) (62), FastTreeMP on XSEDE (2.1.10) (63), and IQTree on XSEDE (2.1.2) (34). MAFFT was used to align the multiple sequences gathered by HOMDscrape and was run at the standard parameters set by CIPRES **(Figure S16)**. To remove unreliable alignment regions, the result of the MAFFT alignment was run through TrimAl at the standard parameters **(Figure S17)**.

### Phylogenetic tree reconstruction

IQTree (2.1.2) was used to create maximum likelihood trees. It was run with the standard parameter settings with 10,000 ultrafast bootstrap replicates to generate branch support statistics **(Table S4)**. FastTree (2.1.10) was used to create secondary fast approximate maximum likelihood trees and was run at the standard parameters set by CIPRES **(Figure S18)**. The constructed trees were then visualized and annotated using iTOL’s online tool (version 6.5.2)(64). A 16S rRNA tree was created using the 16S rRNA RefSeq alignment file (V15.2) available on the HOMD website.

### Tanglegrams for SprA evolution

Tanglegrams were created using Dendroscope v3.8.5 to compare phylogenetic trees of the full SprA and each domain of SprA (40). Species names from the 16S rRNA tree were linked to their respective species name in the phylogenetic trees of SprA and its domains.

### Protein structure alignment

Template sequences for *Capnocytophaga gingivalis* GldJ and GldK were aligned using MAFFT as described above. Structures of GldJ and GldK without signal peptides were generated by Alphafold2 with MMseqs2 (ColabFold v1.5.2) (49, 65). The structures were aligned and visualized using PyMol (66) alignment tool with align method that uses five iteration cycles and a cutoff of 2 Å.

## DECLARATIONS

### Ethics approval and consent to participate

Not applicable.

### Consent for publication

Not applicable.

### Data availability

A version of HOMDscrape is freely available on Github and licensed with an MIT license (https://github.com/strocha1/HOMDscrape). All data generated or analyzed during this study are available in this article and its supplementary files.

## Author Contributions

STR performed majority of the analysis and developed HOMDscrape, DDS performed the protein structure analysis, QZ provided advice on the phylogenetics analysis, STR and AS conceptualized the project and wrote the paper.

## Acknowledgements

We thank Floyd E. Dewhirst for helpful suggestions.

## Supplementary Information

### Supplementary Figures

**Figure S1.**
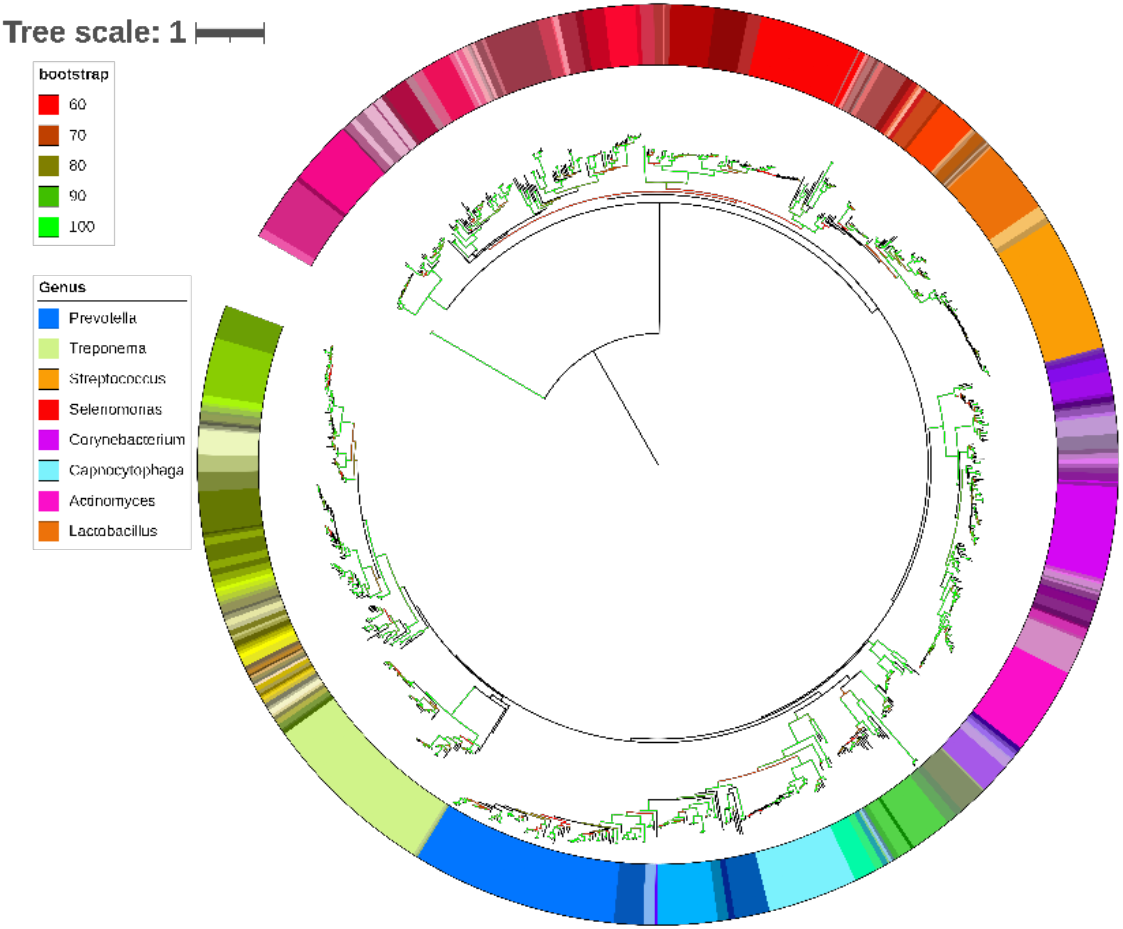
IQ tree of 16S rRNA RefSeq from eHOMD.

**Figure S2.**
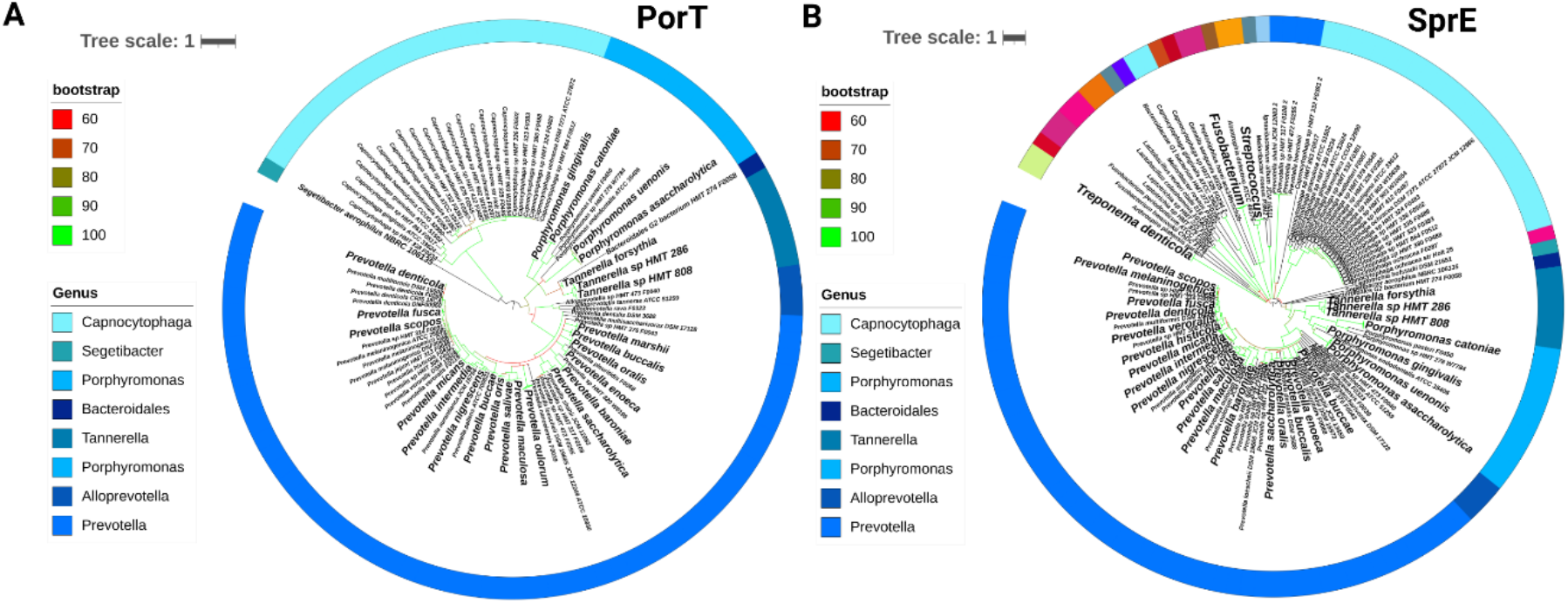
A. IQ Tree of PorT, B. IQ Tree of SprE.

**Figure S3.**
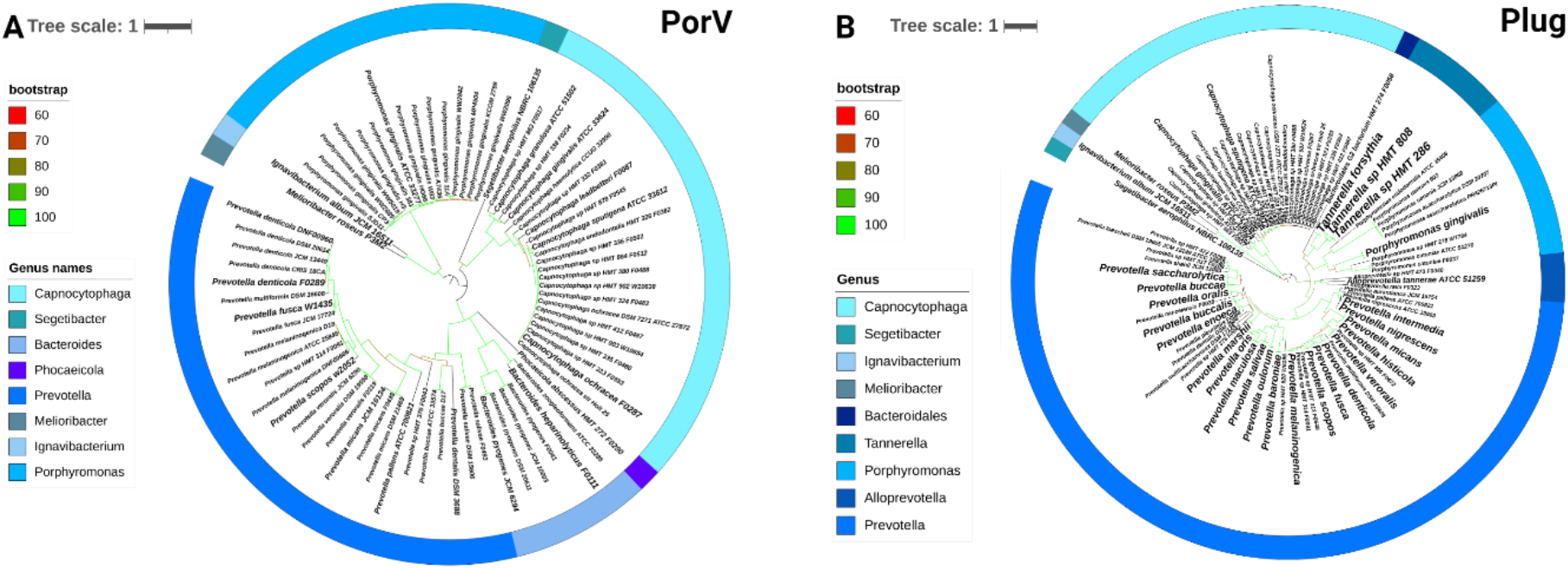
A. IQ Tree of PorV, B. IQ Tree of the Plug.

**Figure S4.**
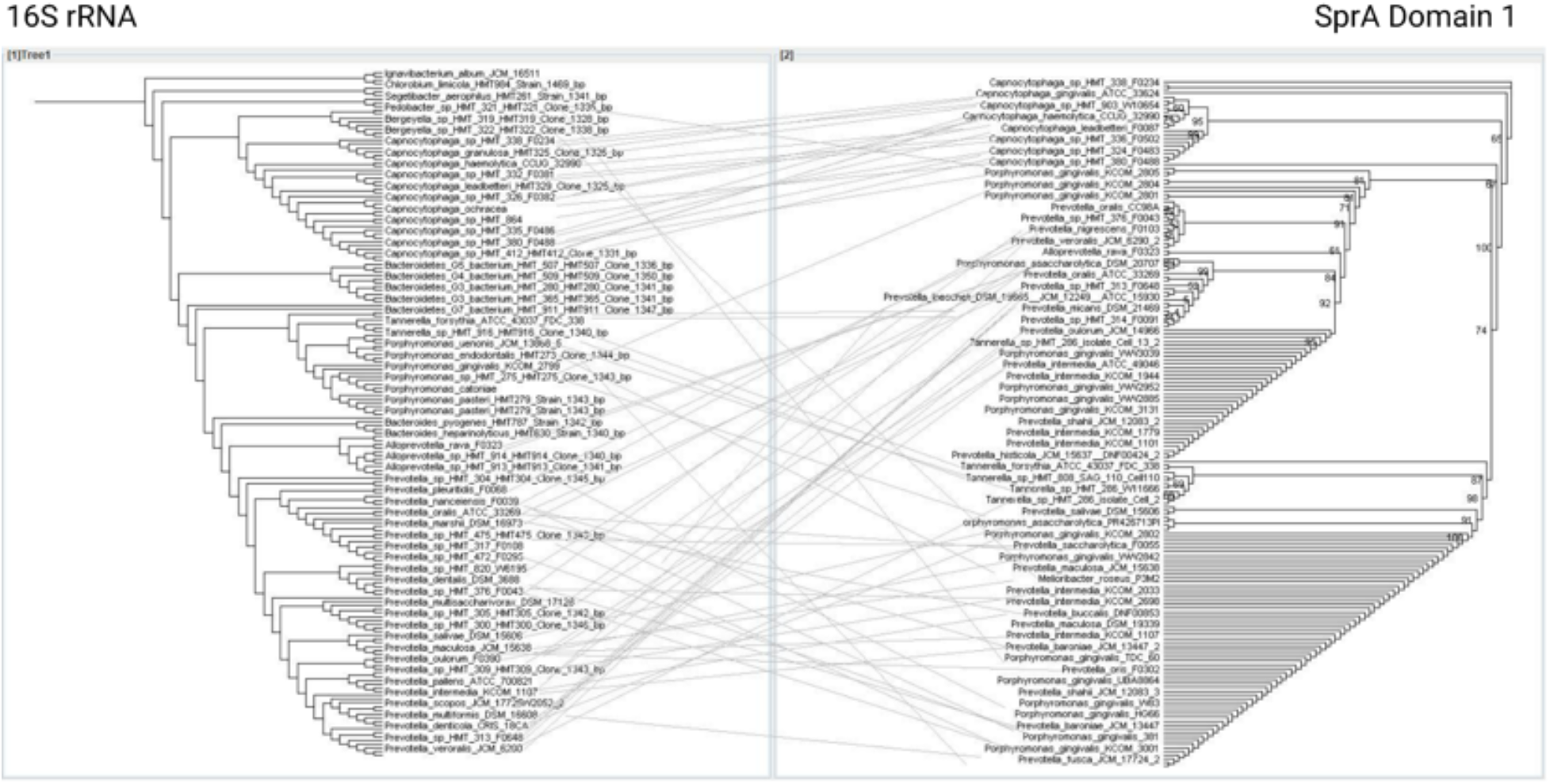
Tanglegram of 16S rRNA tree (Left) and the first domain of SprA (Right)

**Figure S5.**
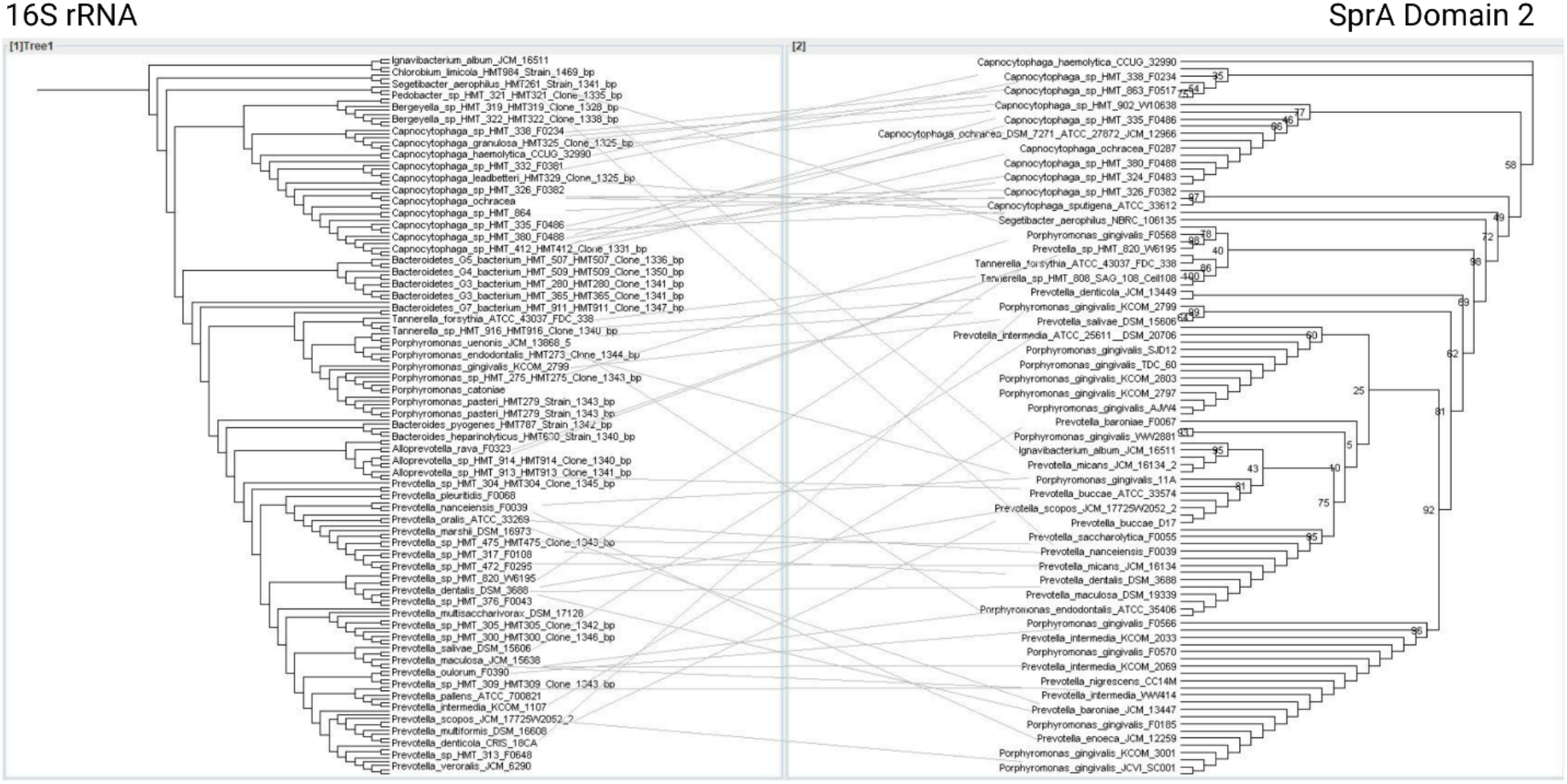
Tanglegram of 16S rRNA tree (Left) and the second domain of SprA (Right)

**Figure S6.**
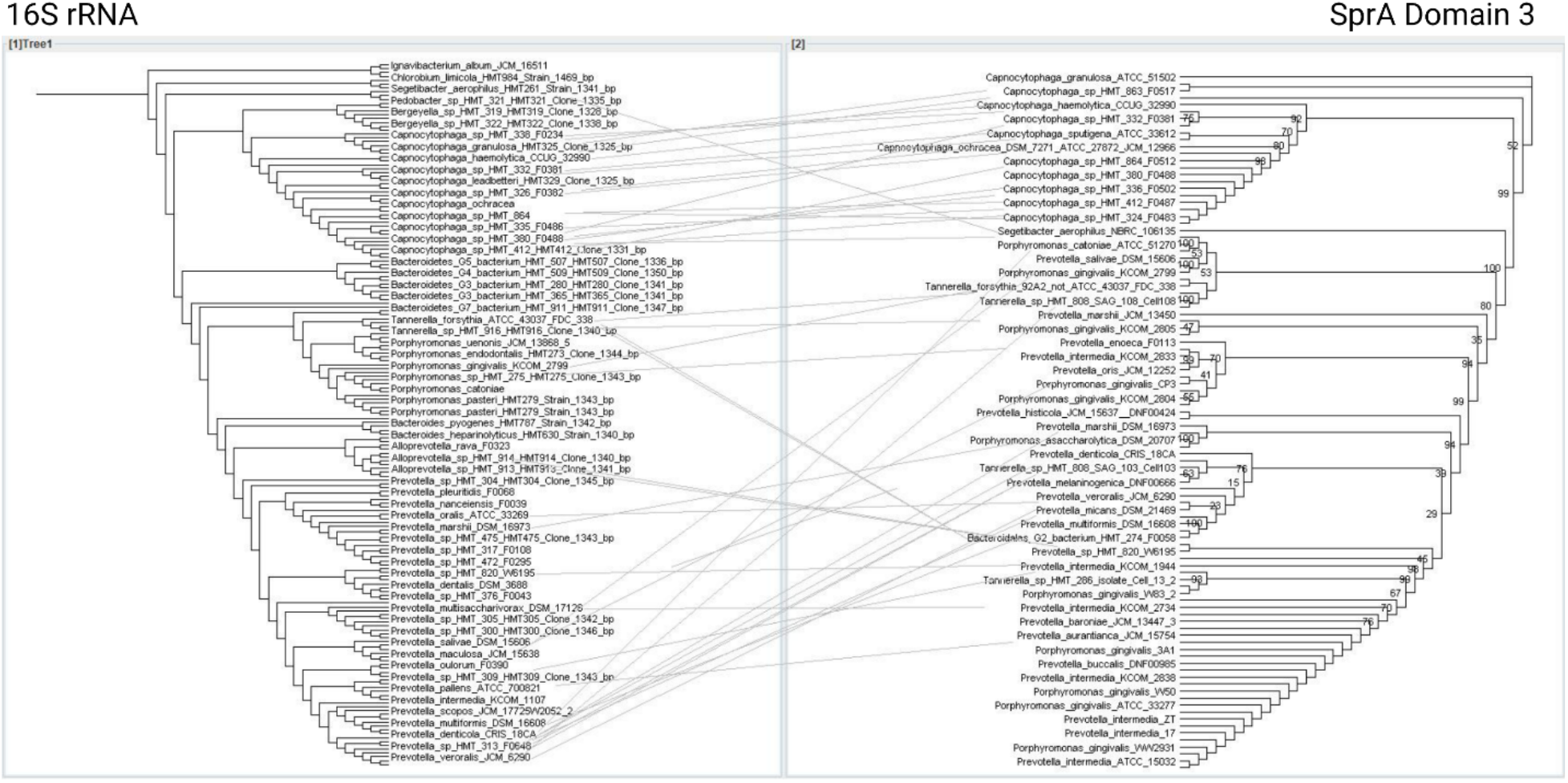
Tanglegram of 16S rRNA tree (Left) and the third domain of SprA (Right)

**Figure S7.**
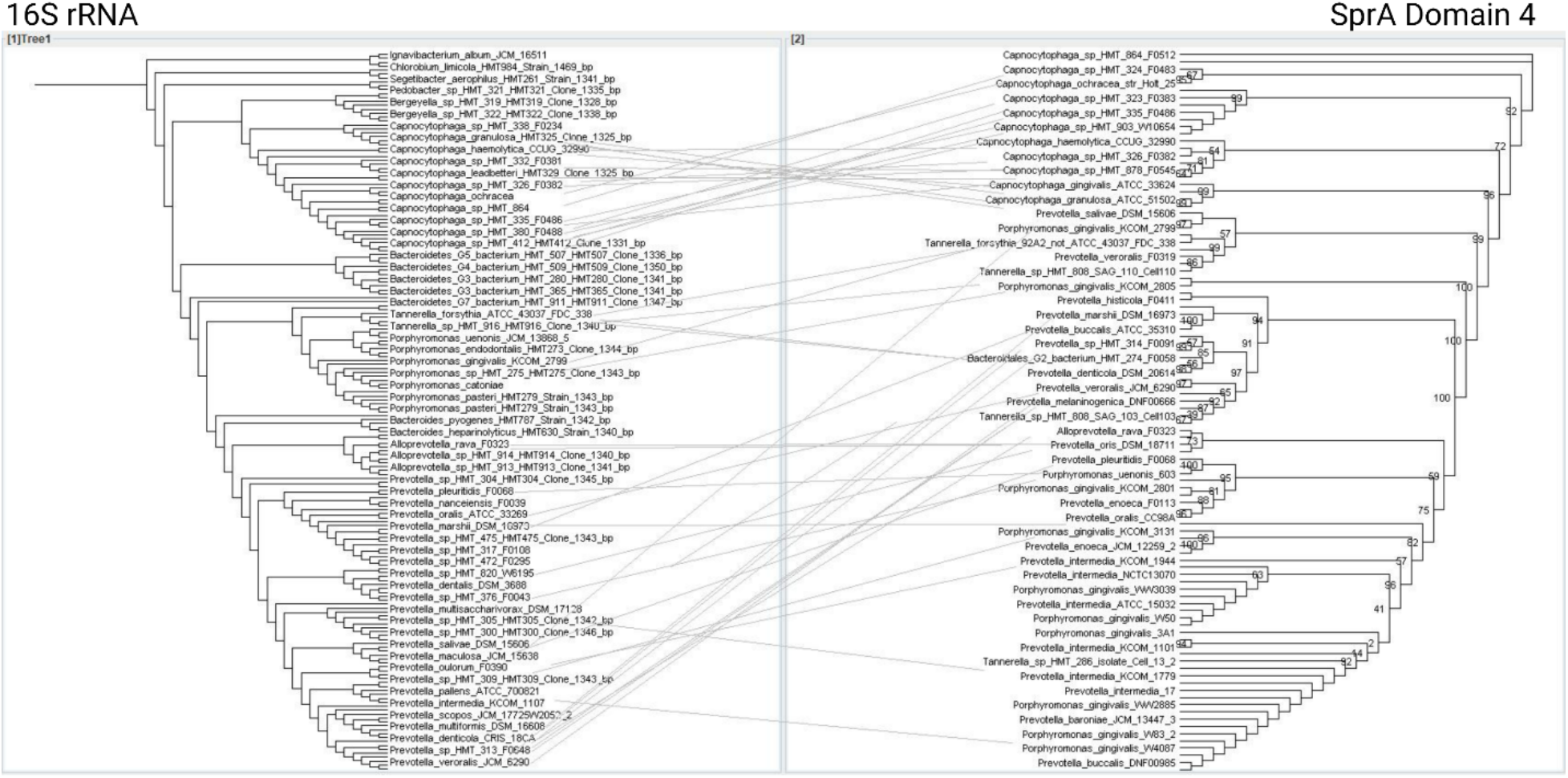
Tanglegram of 16S rRNA tree (Left) and the fourth domain of SprA (Right)

**Figure S8.**
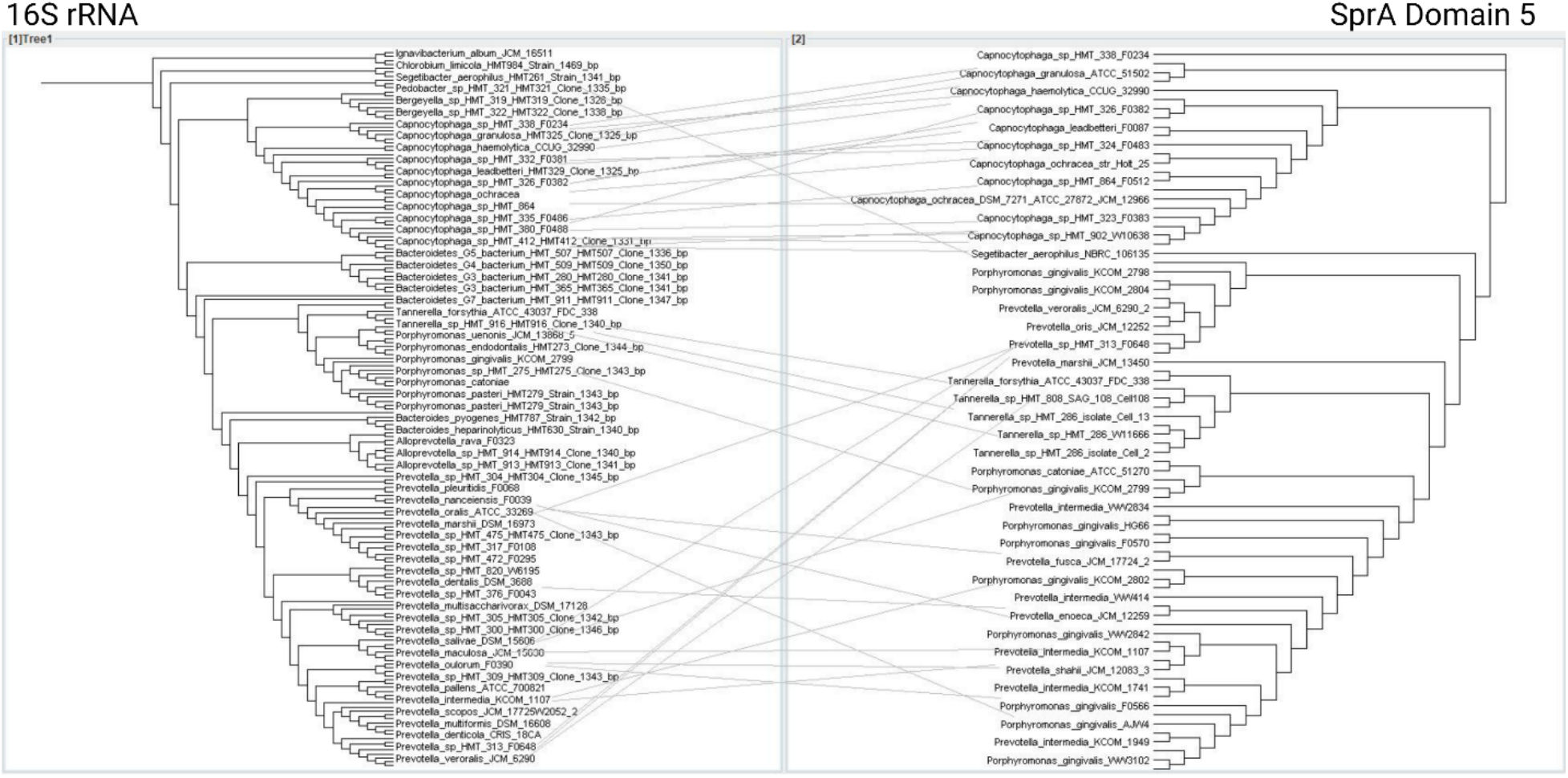
Tanglegram of 16S rRNA tree (Left) and the fifth domain of SprA (Right)

**Figure S9.**
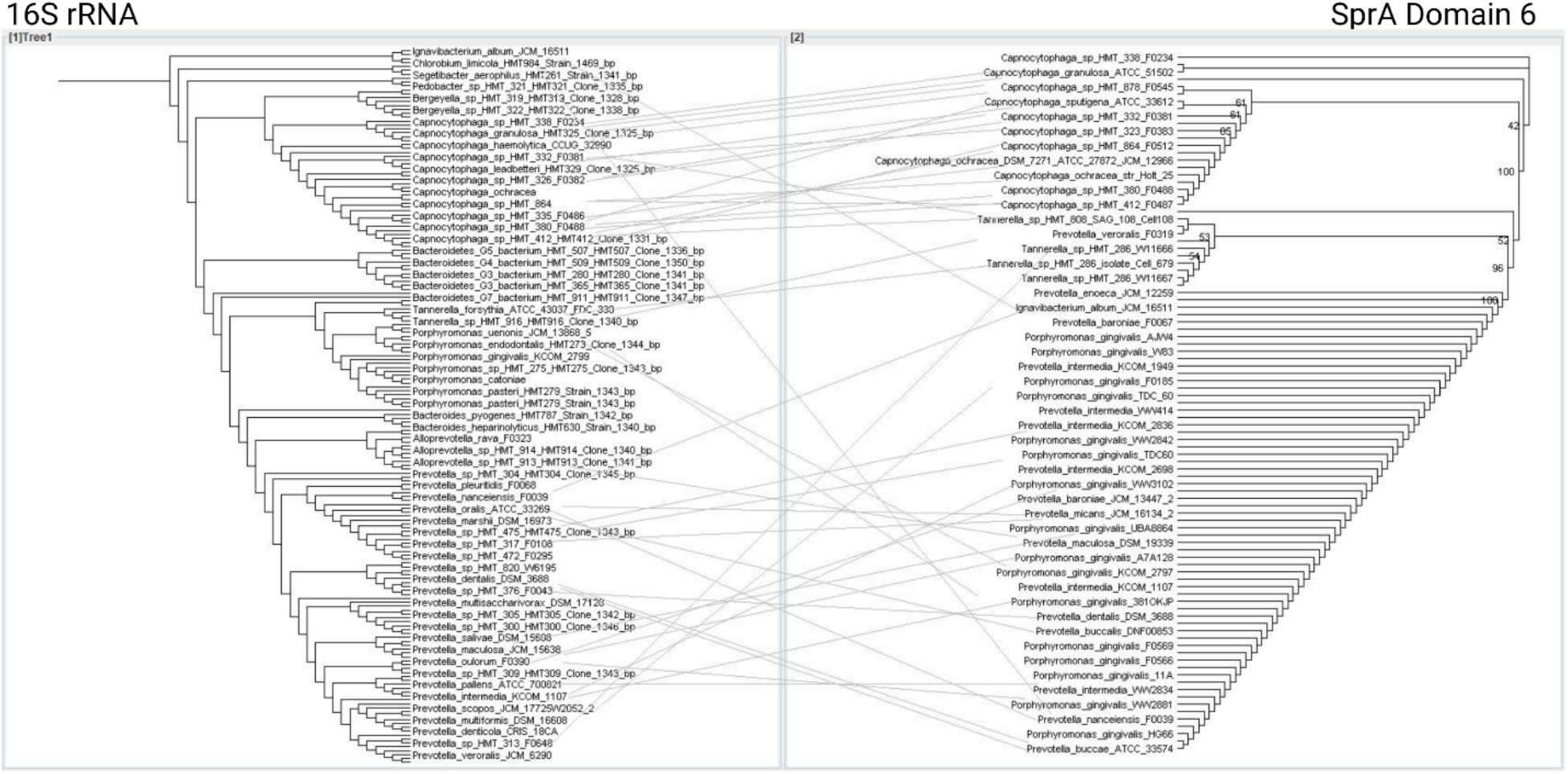
Tanglegram of 16S rRNA tree (Left) and the sixth domain of SprA (Right)

**Figure S10.**
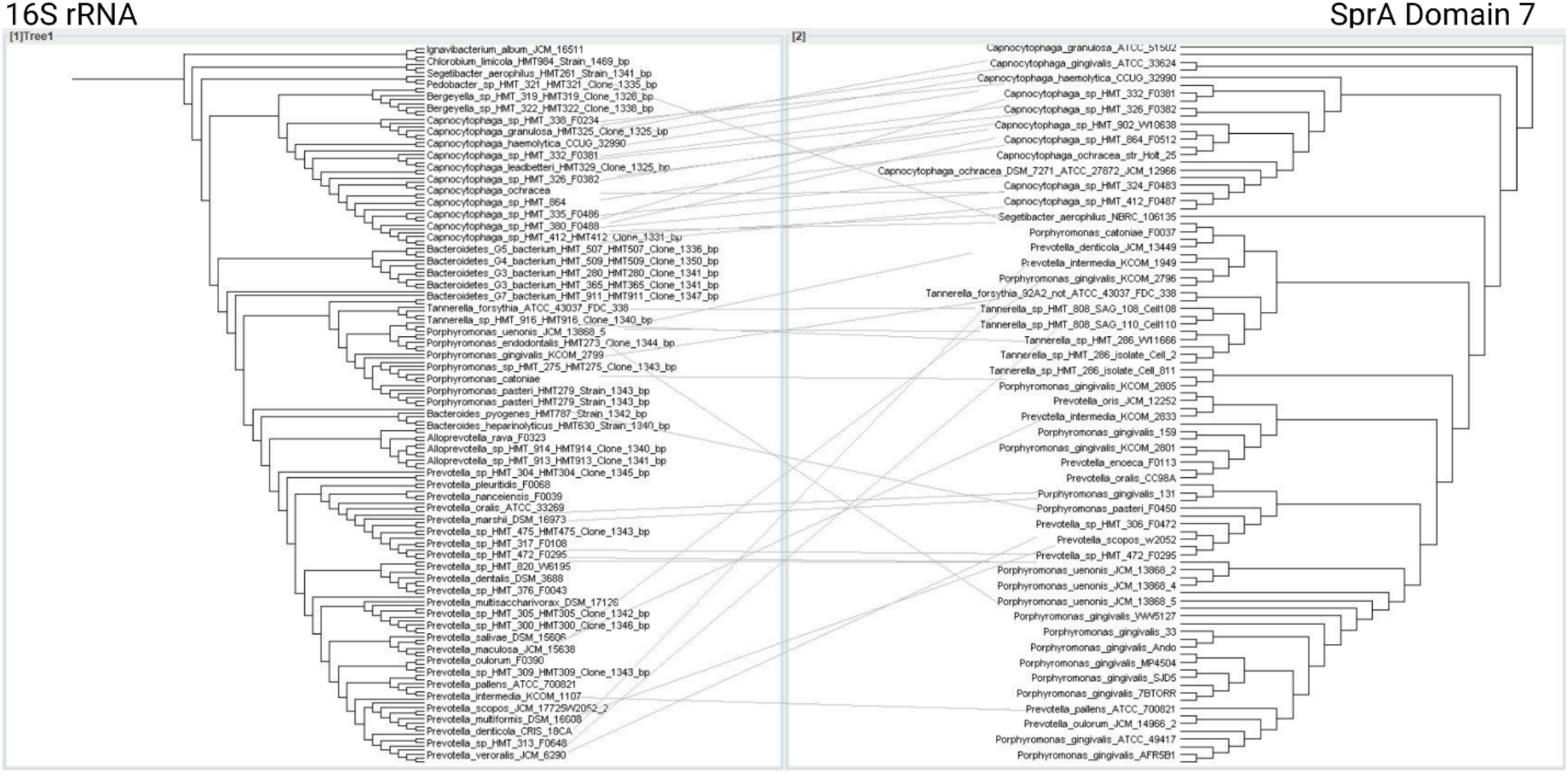
Tanglegram of 16S rRNA tree (Left) and the seventh domain of SprA (Right)

**Figure S11.**
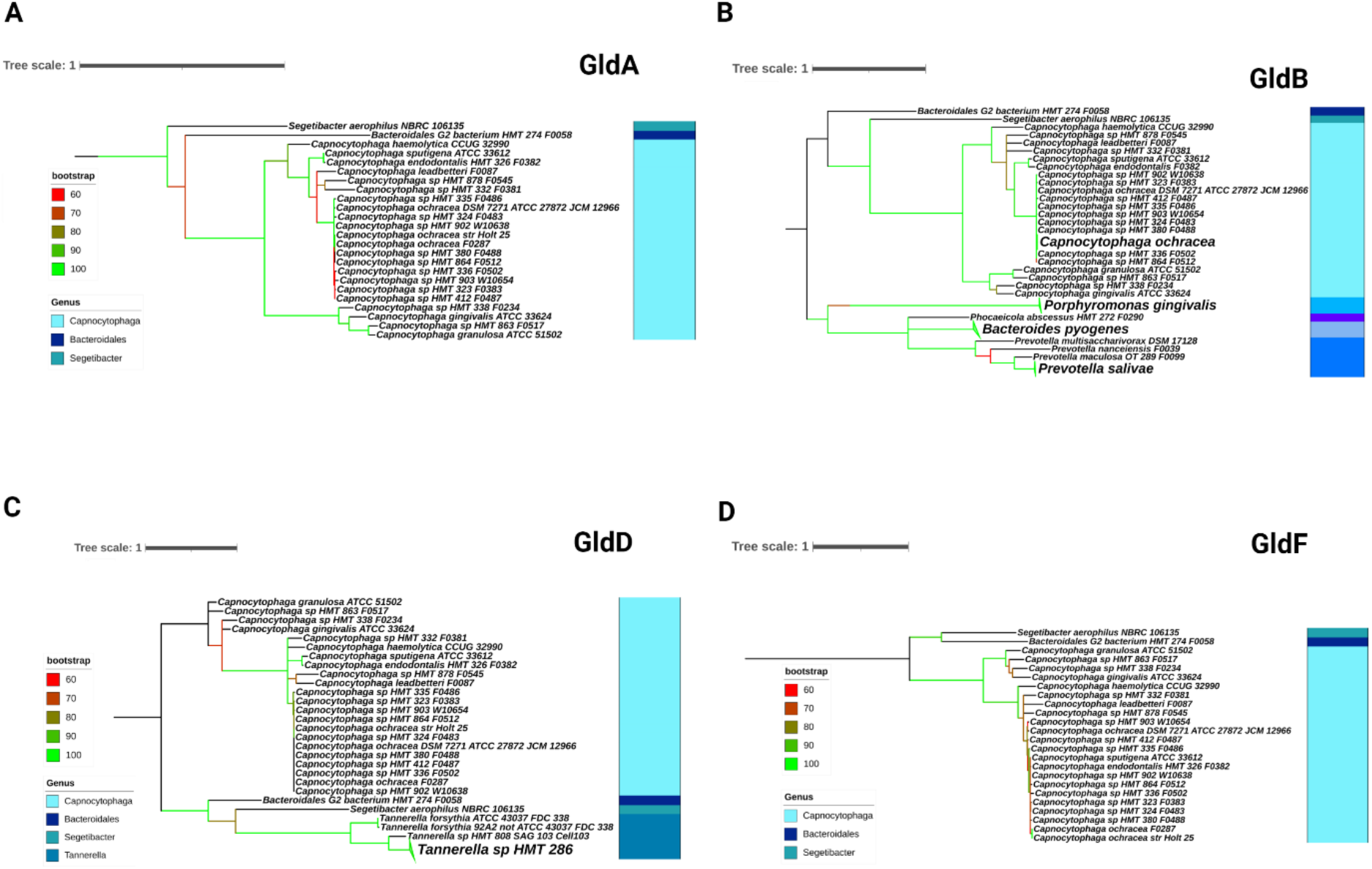
A. IQ Tree of GldA, B. IQ Tree of GldB, C. IQ Tree of GldD, D. IQ Tree of GldF.

**Figure S12.**
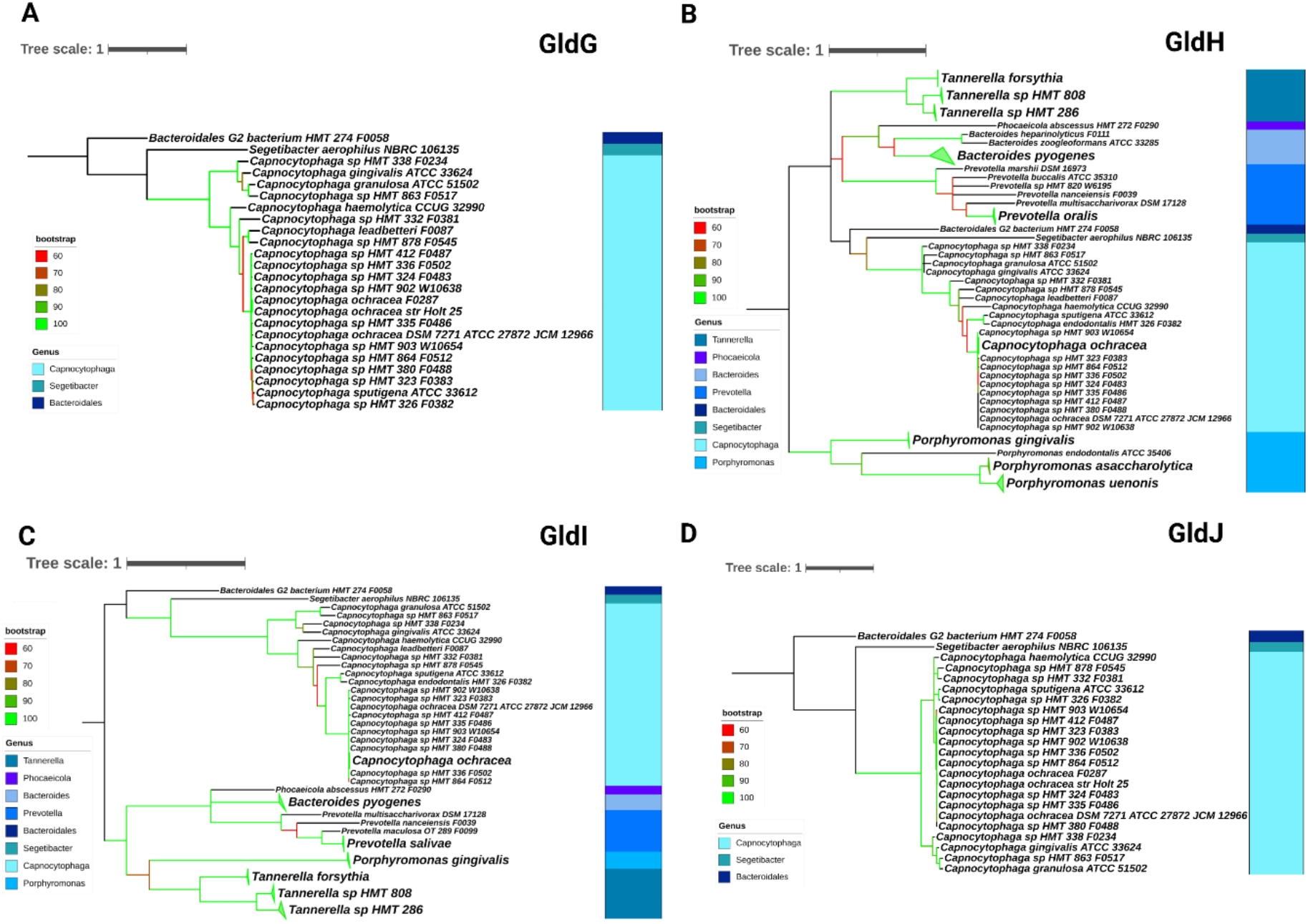
A. IQ Tree of GldG, B. IQ Tree of GldH, C. IQ Tree of GldI, D. IQ Tree of GldJ.

**Figure S13.**
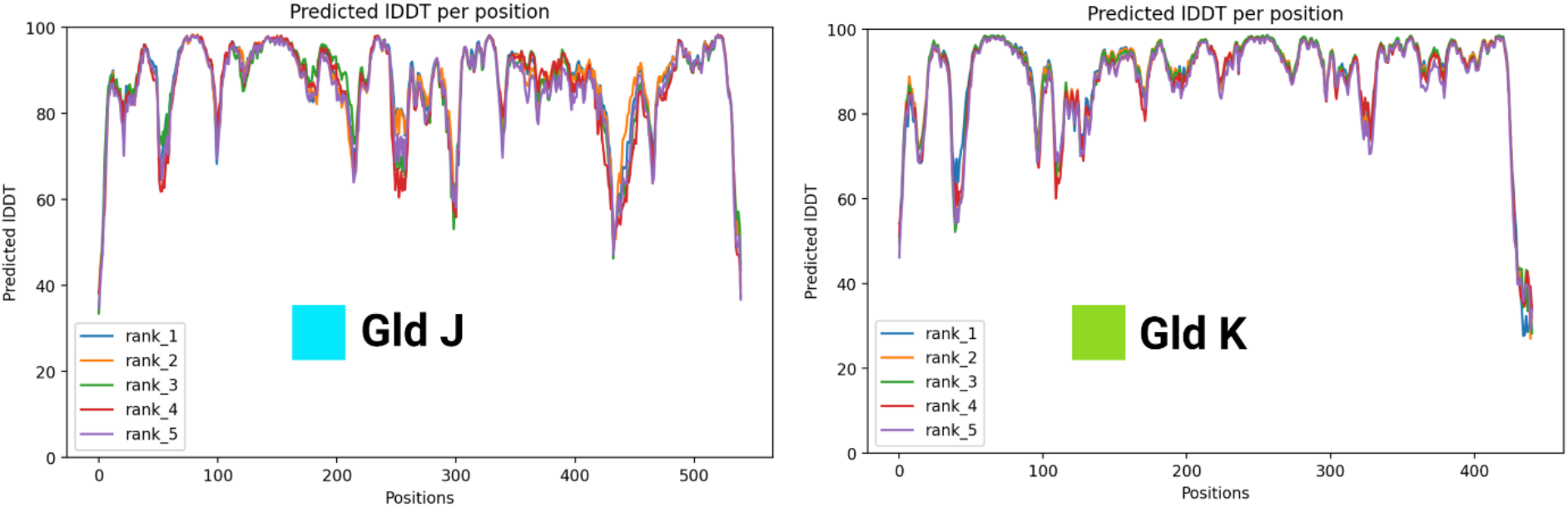
Graphs of the local distance difference test score vs. residue numbers for CG GldJ and CG GldK.

**Figure S14.**
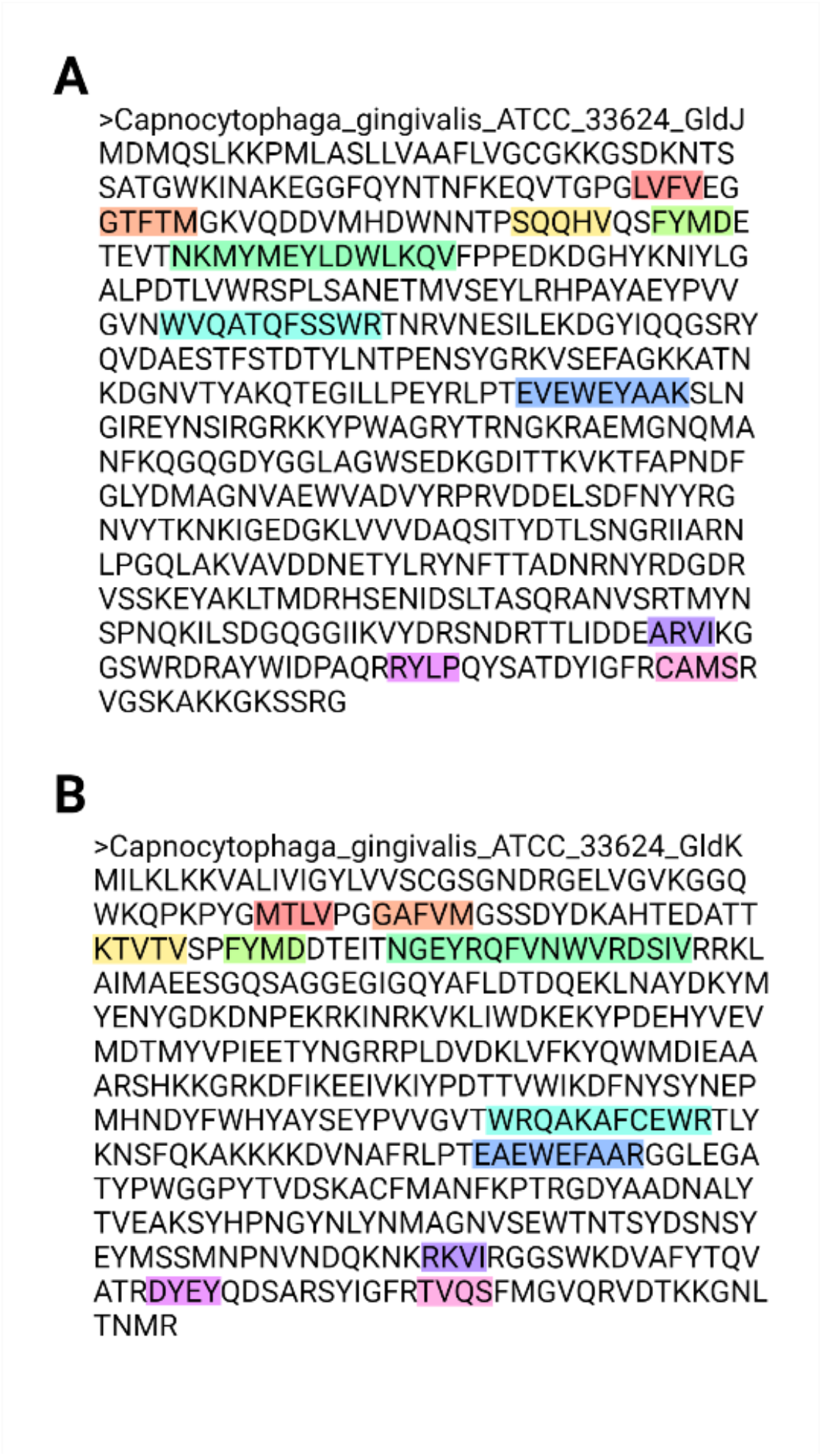
A. Amino acid sequence for GldJ. B. Amino acid sequence for GldK. Regions for structure similarity are highlighted. Each color corresponds to a separate area of similarity.

**Figure S15.**
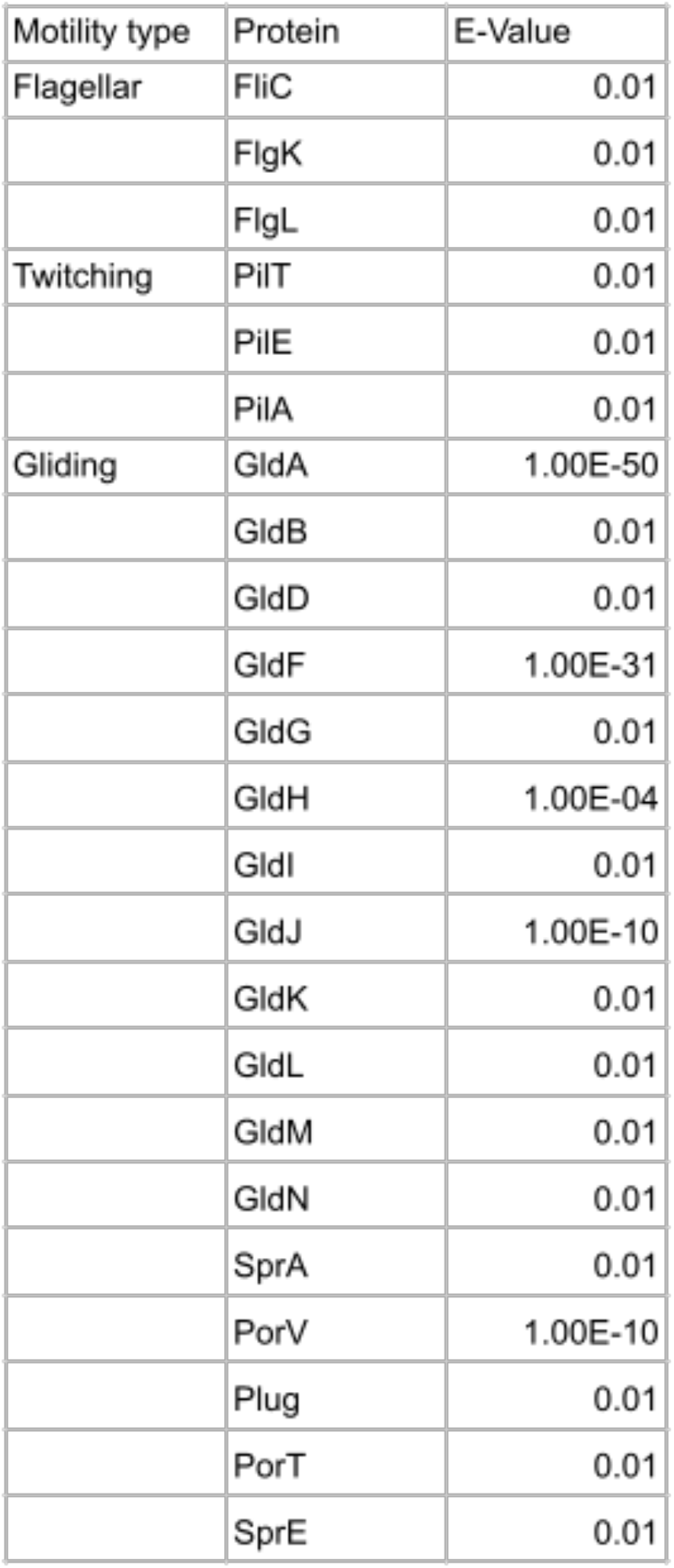
Table describing the E-values used for HOMD BLAST.

**Figure S16.**
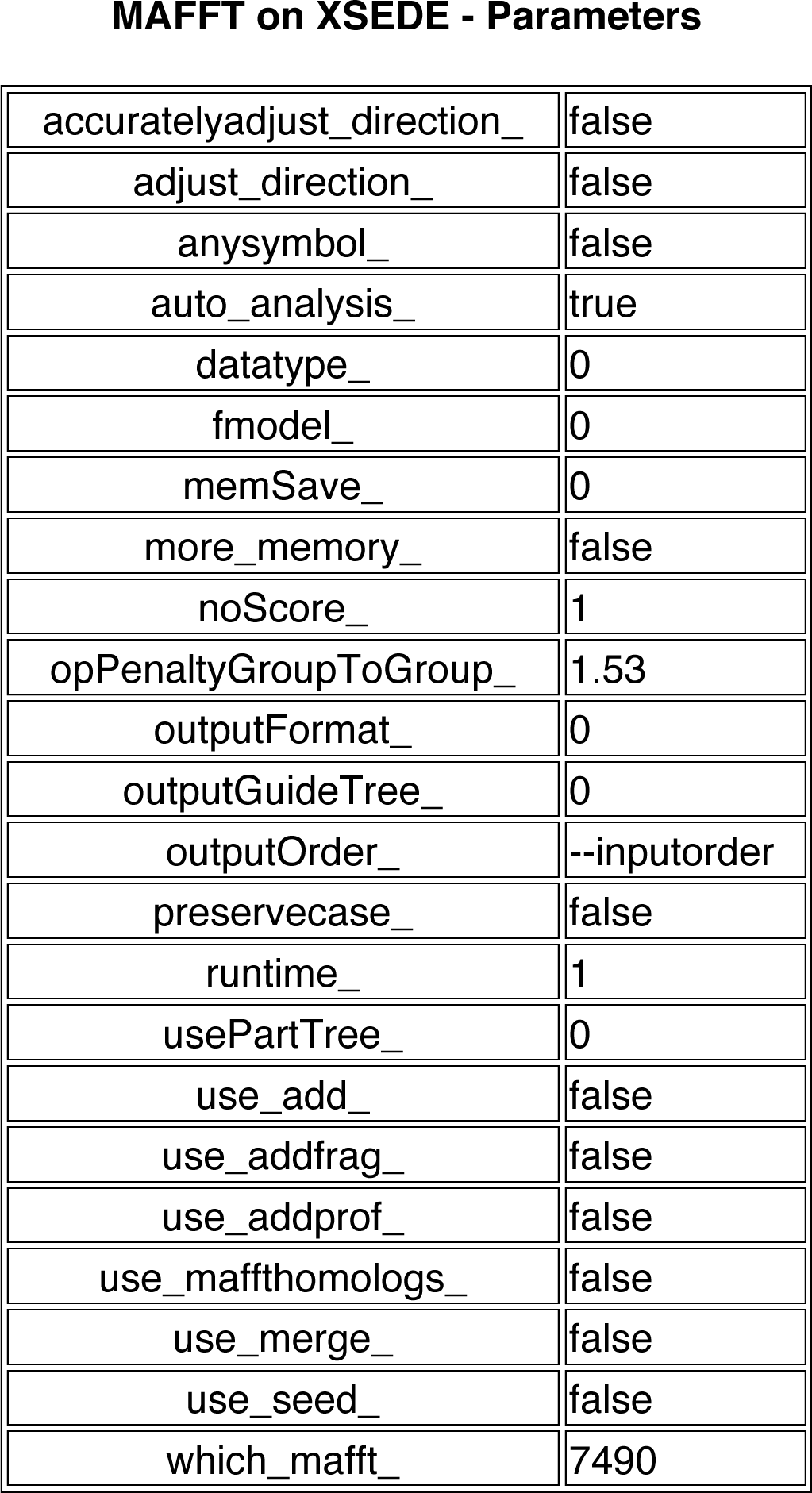
Parameters set for all MAFFT files.

**Figure S17.**
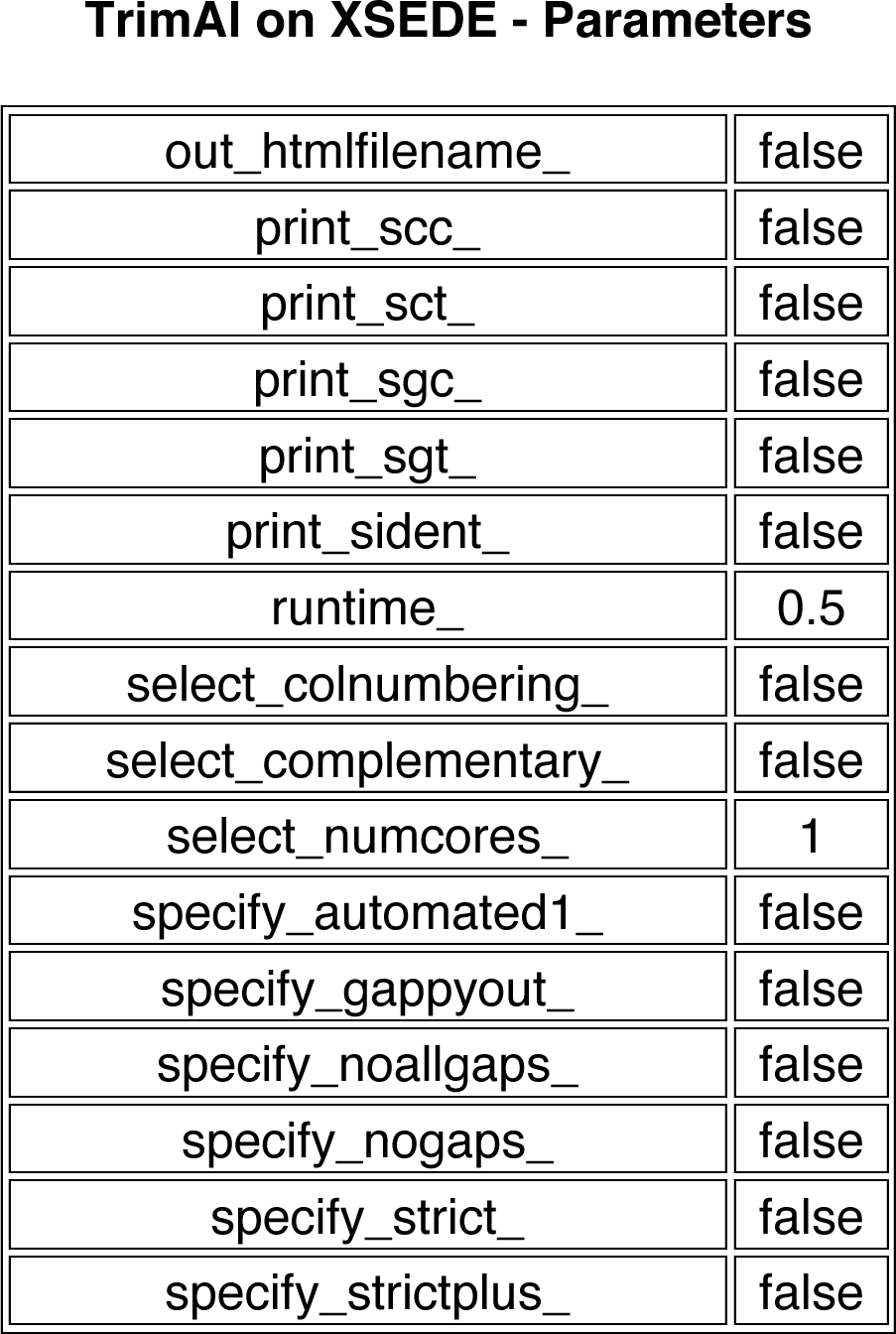
Parameters set for all TrimAl files.

**Figure S18.**
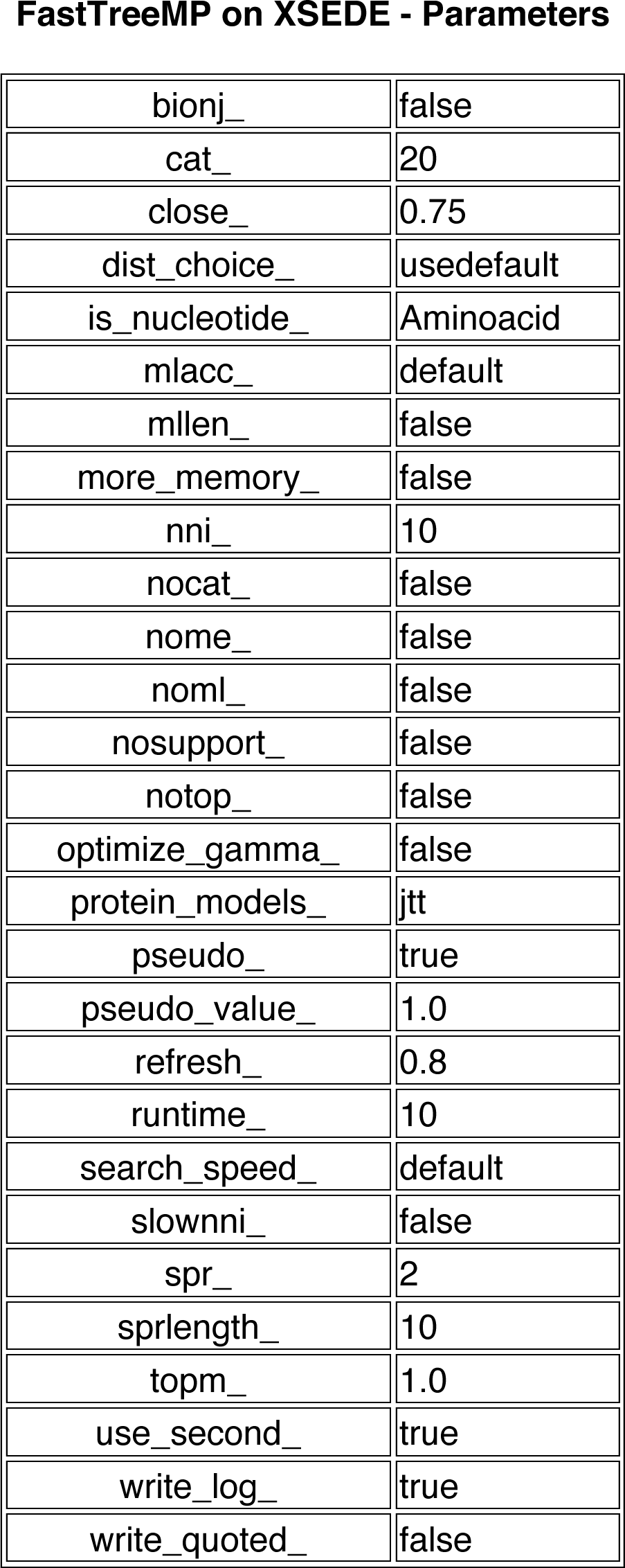
Parameters set for all FastTreeMP files.

### Supplementary Tables

**Table S1.** Comparison table of the flagellar motor and the virulence-associated type III injectosome structural proteins.

**Table S2.** Summary of all species found on HOMD and their links to motility.

**Table S3.** Comparison table of type 4 pilus driven twitching motility proteins and type 2 secretion system proteins.

**Table S4.** Parameters for IQ-Tree

